# Spatiotemporal organization of movement-invariant and movement-specific signaling in the output layer of motor cortex

**DOI:** 10.1101/2020.10.27.357087

**Authors:** Stephen P. Currie, Julian J. Ammer, Brian Premchand, Joshua Dacre, Yufei Wu, Constantinos Eleftheriou, Matt Colligan, Thomas Clarke, Leah Mitchell, Aldo Faisal, Matthias H. Hennig, Ian Duguid

## Abstract

Motor cortex generates descending output necessary for executing a wide range of limb movements. Although movement-related activity has been described throughout motor cortex, the spatiotemporal organization of movement-specific signaling in deep layers remains largely unknown. Here, we recorded layer 5B population dynamics in the caudal forelimb area of motor cortex while mice performed a forelimb push/pull task and found that most neurons show movement-invariant responses, with a minority displaying movement specificity. Cell-type-specific imaging identified that movement-invariant responses dominated pyramidal tract (PT) neuron activity, with a small subpopulation representing movement type, whereas a larger proportion of intratelencephalic (IT) neurons displayed movement-specific signaling. The proportion of IT neurons decoding movement-type peaked prior to movement initiation, while for PT neurons this occurred during movement execution. Our data suggest that layer 5B population dynamics largely reflect movement-invariant signaling, with information related to movement-type being differentially routed through relatively small, distributed subpopulations of projection neurons.

## Introduction

In mammals, descending cortical output is necessary for the learning and execution of voluntary movements (Guo et al., 2015; Hwang et al., 2019; Hwang et al., 2021; Kawai et al., 2015; Lawrence and Kuypers, 1968; Martin and Ghez, 1991). Deep layer projections from primary motor cortex form multiple descending pathways innervating cortical, subcortical, brainstem and spinal cord circuits necessary for triggering and controlling movement (for review, see Lemon, 2008; Ruder and Arber, 2019; Shepherd, 2013). Individual layer 5 projection neurons display complex firing patterns that correlate with various aspects of limb trajectories, such as joint angle, direction, and speed (Georgopoulos et al., 1982; Moran and Schwartz, 1999; Paninski et al., 2004; Thach, 1978) and during single action tasks in rodents, most layer 5 projection neurons (>70%) display movement-related activity in the form of bidirectional changes in firing rate (Dacre et al., 2021; Estebanez et al., 2017; Levy et al., 2020; Park et al., 2021; Sauerbrei et al., 2019; Wang et al., 2017), suggesting widespread encoding of movement. However, in non-human primates the largest components of motor cortex population responses during a delayed-multi-direction reach task have been shown to be “condition-invariant”, meaning the population response magnitude and time course were similar irrespective of reach direction (Kaufman et al., 2016). Condition-invariant responses are tightly linked to the onset of movement likely reflecting movement timing rather than movement type, similar to condition-invariant population transitions observed in recurrent neural networks trained to recapitulate complex muscle patterns in reaching primates (Sussillo et al., 2015). Deciphering how condition-invariant (which we term “movement-invariant”) and movement-specific signaling is spatiotemporally organized in the output layers of motor cortex and how they map onto specific projection classes would provide an important step towards understanding descending cortical control of movement.

In rodents, descending information from the main output layer of motor cortex, layer 5B, is routed via two molecularly and anatomically defined projection pathways: pyramidal tract (PT) neurons innervate multiple targets including thalamus, subthalamic nucleus, superior colliculus, ipsilateral striatum, brainstem and spinal cord, but not cortex or contralateral striatum (Economo et al., 2018; Kita and Kita, 2012; Muñoz-Castañeda et al., 2021; Ueta et al., 2014; Winnubst et al., 2019); while intratelencephalic (IT) neurons target bilateral striatum and cortex, but not other subcortical targets (Levesque et al., 1996; Muñoz-Castañeda et al., 2021; Wilson, 1987; Winnubst et al., 2019). Although layer 5B neurons are reciprocally connected (Kiritani et al., 2012; Morishima and Kawaguchi, 2006; Morishima et al., 2011), connectivity is essentially unidirectional from IT to PT neurons (Kiritani et al., 2012). This form of asymmetric across-projection class connectivity appears to be a common cortical motif necessary for sensorimotor processing (Brown and Hestrin, 2009; Kiritani et al., 2012; Molyneaux et al., 2007; Reiner et al., 2010). From a descending control perspective, asymmetric connectivity coupled with differential PT and IT intrinsic excitability, sensitivity to neuromodulation and local- and long-range inputs (for review, see Baker et al., 2018; Shepherd, 2013) provides a mechanism for flexible routing of information via distinct output channels depending on behavioral state and task requirements. Accumulating evidence suggests that PT neurons provide an essential source of descending control signals for the execution of voluntary limb movements (Economo et al., 2018; Nelson et al., 2021; Peters et al., 2017; Soma et al., 2017; Wang et al., 2017), while IT neurons provide input to cortical and striatal circuits contributing to movement preparation and specification (Panigrahi et al., 2015; Park et al., 2021; Yttri and Dudman, 2016), but how movement-specific information is spatiotemporally organized across the two output channels remains unclear.

Here, we performed 2-photon calcium imaging in deep layers of the caudal forelimb area (CFA) of mice trained to perform two diametrically opposing forelimb movements (i.e., an alternating push / pull lever task). By combining population imaging with neural classifiers and dimensionality reduction, we show that the majority of layer 5B neurons display movement-invariant signaling, correlated with movement timing rather than movement type, while small subpopulations of PT and IT neurons convey movement-specific information. Decoding of movement type was most prevalent prior to movement initiation in IT neurons, and during movement execution in PT neurons, with high decoding accuracy neurons from both projection classes being temporally uncorrelated and distributed across layer 5B. These findings provide evidence that movement-invariant signaling dominates layer 5B activity, while movement-specific information is spatially and temporally distributed across molecularly and anatomically defined projection neuron classes.

## Results

### CFA is required for execution of a cued push / pull lever task for mice

To explore how layer 5B signaling relates to the execution of different movements, we first developed a cued linear push / pull lever task for mice. The task design required mice to push or pull a horizontal lever during presentation of a 2 second 6 kHz auditory cue in order to receive a water reward. After a 4-6 s inter-trial interval (ITI), mice had to push the lever 4 mm forward from an initial starting position, the lever would then be locked, and a servo-controlled lick spout would deliver a 5 μl reward following a 1s delay. The lever was then unlocked and a second ITI commenced where mice would then be expected to pull the lever backwards 4 mm to the original starting position after presentation of the same 6 kHz auditory cue. Miss trials or spontaneous movements during the ITI resulted in a lever reset and restarting of the ITI (Figure 1A). Individual mice displayed idiosyncratic strategies to engage with the lever but displayed reproducible trial-to-trial forelimb trajectories (Figure 1B). In general, mice reoriented their forelimb and paw upon cue presentation (lift and rotate backwards for pushes, lift and rotate forwards for pulls) during movement initiation (Video S1). Mice rapidly learned the task (mean 10.5 days, [4] inter-quartile range (IQR), N = 24 mice), displaying fast reaction times and movement durations that reflect the combination of paw reorientation and lever manipulation (Figure 1C). ‘Expert’ mice completed 44.5 [9.5] IQR successful push and 45.0 [8.5] IQR successful pull trials during each 30-minute behavioral session equating to ∼71 % task success (push median 68.0 % [35.9] interquartile range (IQR); pull median 74.5 % [43.4] IQR; N = 24 mice) (Figure 1D-1E).

**Figure 1.**
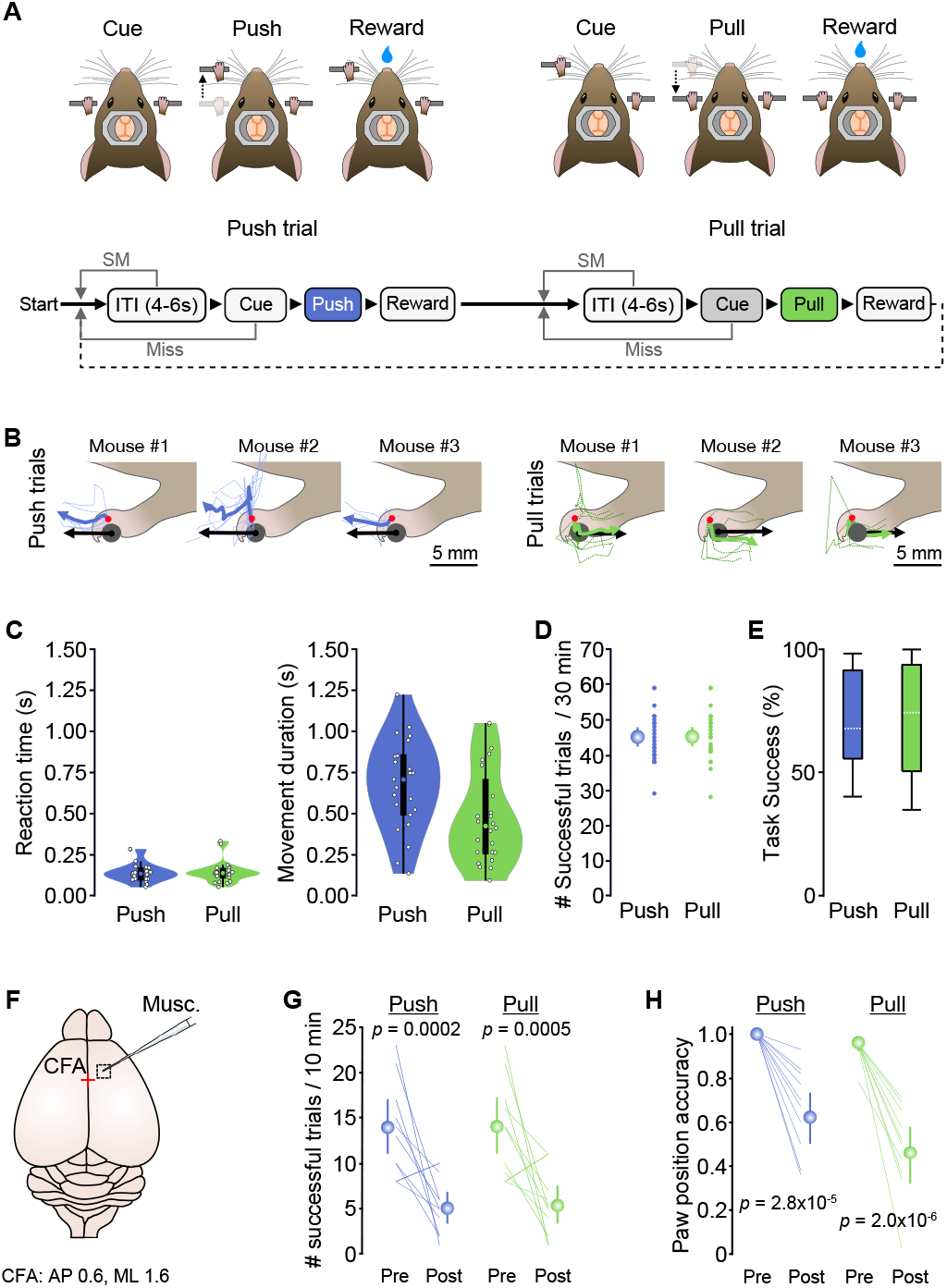
Caudal forelimb area is necessary for executing cued push and pull movements. (A) *Top*, cued alternating push / pull task for head restrained mice. Bottom, behavioral task structure: ITI, inter-trial interval; SM, spontaneous movement. (B) Paw and lever movement trajectories from 3 mice presented relative to position at movement initiation. Single trial (dashed lines) and mean paw trajectories (solid lines) during push (left; blue) and pull (right; green) trials are shown alongside the average movement vector of the lever (black arrow). Red dot, depict approximate tracked position on the paw at movement initiation. (C) Violin plots showing median, interquartile range and range of reaction times (left) and movement durations (right) during push (blue) and pull (green) trials. Circles represent data from individual mice (N = 24 mice). (D) Number of successful trials per 30 minute training session (small symbols, data from individual mice; large symbols, mean ± 95% CI; N = 24 mice). (E) Box-and-whisker plots showing median, interquartile range and range of task success across mice (N = 24 mice). (F) Focal muscimol inactivation of CFA, centred on 0.6 mm anterior, 1.6 mm medial of bregma. Red cross denotes bregma. (G) Number of successful push (blue) and pull (green) trials in a 10 min period before (Pre) and after (Post) injection of muscimol (N = 10 mice), paired t test. Colored lines, individual mice. Symbols, population means ± 95% CI. (H) Paw position accuracy at the point of cue presentation before (Pre) and 10 mins after (Post) muscimol injection into CFA (N = 10 mice), paired t test. Colored lines, individual mice. Symbols, population means ± 95% CI.

To confirm that the caudal forelimb area (CFA) of motor cortex is required for the execution of both push and pull movements, we focally injected the GABA_A_ receptor agonist muscimol (1.6 mm lateral, 0.6 mm rostral of bregma, see Methods) (Dacre et al., 2021; Schiemann et al., 2015). By applying muscimol during behavioral engagement, we could assess the immediate effects of CFA inactivation within the first 10 mins following drug injection (Figure 1F), where drug diffusion remained within the targeted region (Figure S1A-S1C). Muscimol rapidly blocked initiation of both actions, reducing the number of successful trials in the first 10 minutes by ∼65% (push Pre 13.9 [11.1 17.1] 95% CI trials, push Post 5.0 [3.3 6.8] 95% CI trials, N = 10 mice; pull Pre 14.0 [10.9 17.2] 95% CI trials, pull Post 5.3 [3.2 7.4] 95% CI trials, N = 10 mice) (Figure 1G). Sham injections of saline into CFA or muscimol injections into hindlimb motor cortex had no effect (Figure S1D-1G). Blocking activity in CFA resulted in both an inability to initiate push or pull movements, and monoparesis of the contralateral forelimb (i.e., localized weakness without complete loss of function), as evidenced by the significant reduction in paw position accuracy (i.e., the forepaw was not positioned on the lever at cue presentation) (Figure 1H, Figure S1H-1J and Video S2). The effect of muscimol inactivation was most pronounced in mice that displayed the highest number of successful trials, confirming that task execution is CFA-dependent even in expert mice (Hwang et al., 2019; Hwang et al., 2021; Kawai et al., 2015) (Figure S1K).

### Movement-invariant signaling dominates layer 5B activity patterns

To examine how output from CFA relates to the execution of push and pull movements, we restricted imaging of behavior-related population activity to cortical depths corresponding to layer 5B, the upper boundary of which was identified by the presence of pyramidal tract neurons in separate tracing experiments (boundary ≥ 500 μm from the pial surface at the center of CFA) (Schiemann et al., 2015) (Figure 2A-2B and Figure S2A). Cell density estimates suggested that we imaged the majority of layer 5B neurons at depths up to 650 μm from the pial surface (Figure S2A-C). A large proportion of layer 5B neurons displayed movement-related activity (468/653 neurons, mean = 73.5% [54.7 81.8] 95% CI per field-of-view (FOV) (n = 12 FOVs, N = 6 mice), where ΔF/F_0_ changes occurred within a peri-movement window spanning 150 ms prior to movement initiation to 40 ms after median movement completion. The remaining neurons were classified as either non-responsive, or ‘reward-related’ if changes in ΔF/F_0_ occurred after the peri-movement window (Figure 2C-2D). The trial-to-trial similarity in population responses of movement-related neurons strongly correlated with the similarity in forelimb movement magnitude (i.e., motion index) during both push and pull movements, suggesting ΔF/F_0_ changes reflected movement of the forelimb (Figure 2E). By comparing push and pull trials, we found that most layer 5B neurons displayed movement-related activity that was indistinguishable between trial type (median = 59.8 % [31.4] IQR, N = 6), termed ‘movement-invariant’ signaling, manifest as either increased (85 %) or decreased (15 %) activity correlating with the onset of movement. Movement-invariant neurons appeared to reflect the timing of movement (i.e., transition from resting posture to push/pull) rather than movement type and were spatially distributed across each FOV (Figure 2F-2H). In contrast, only a small fraction of neurons displayed movement-specific signaling, where ΔF/F_0_ changes were significantly different between push and pull trials (termed movement bias, push bias - median = 14.3 % [15.9] IQR; pull bias - median = 11.8 % [19.5] IQR, N = 6 mice) (Figure 2M). Responses of movement bias neurons were classified into four different types including both positive and negative changes in ΔF/F_0_, consistent with bidirectional movement-specific tuning of neural activity (Georgopoulos et al., 1982). Most movement-bias neurons were classified as Type 1 (136 / 181 neurons, 75.1 %), showing increased ΔF/F_0_ during both push and pull trials, while smaller proportions of Type 2 (25 / 181 neurons, 13.8 %) and Type 3 (15 / 181 neurons, 8.3 %) neurons displayed movement selectivity (i.e., significant change in ΔF/F_0_ for one movement, with no response during the opposing movement). Finally, a small minority of cells displayed reduced activity during both push and pull trials classified as Type 4 neurons (5 / 181 neurons, 2.8 %, N = 6 mice) (Figure 2I-2J). In terms of spatial organization, movement bias neurons were found in all FOVs and were spatially intermingled with movement-invariant neurons (Figure 2H and 2K). Although there was a high degree of variability in ΔF/F_0_ changes trial-to-trial, no consistent differences in mean pairwise trial-to-trial ΔF/F_0_ correlations were found between movement-invariant and movement-bias neurons across trial types (Figure 2L). A small proportion of bias neurons displayed differences in baseline ΔF/F_0_ between push and pull trials, which could reflect postural differences (i.e., different start positions for push and pull trials) or differential preparatory activity (Li et al., 2015) (Figure S2D-S2F). However, baseline differences were on average smaller than those observed during the peri-movement epoch (data not shown), thus unlikely to be the main driver of movement-specific signaling. Given that ΔF/F_0_ changes provide an indirect readout of neural activity, we sought to confirm the proportions of movement-invariant and movement-bias neurons in layer 5B using high-density silicone probe recordings. Putative layer 5B projection neurons were identified using spike-width thresholding and electrode depth profiling based on retrograde labeling from the pons (Figure S2G-S2I). We found similar proportions of movement-invariant (no bias) and push- / pull-bias neurons when comparing both recording methods (Figure 2M and Figure S2J-S2L), confirming that movement-invariant signaling dominated layer 5B responses, while a small proportion of neurons conveyed movement-specific information.

**Figure 2.**
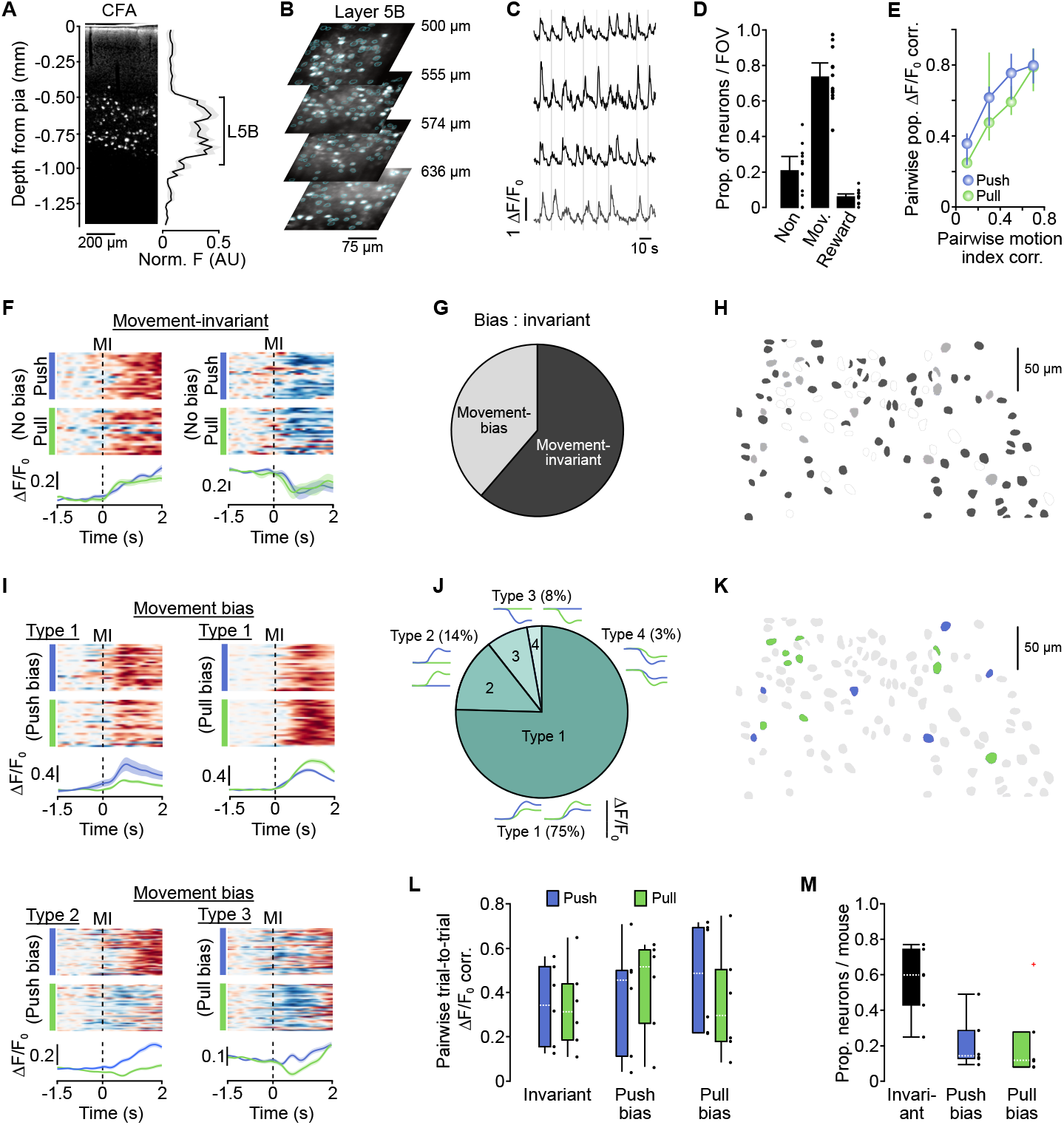
Movement-invariant and movement-specific signaling in layer 5B of CFA. (A) Depth profile of pyramidal tract (PT) neurons in CFA: Left, Retrobead labeling of PT neurons following injection into the pons; Right, normalized fluorescence ± s.e.m as a function of depth from the pial surface (N = 3 mice). Black line indicates the upper and lower boundary of layer 5B across mice. (B) Representative 2-photon imaging fields-of-view (FOVs) from layer 5B in CFA (N = 4 mice). Cyan circles depict regions of interest. (C) ΔF/F_0_ traces from four example layer 5B neurons during task execution (gray vertical bars). Black traces depict neurons with movement-related activity; gray trace depicts a neuron with reward-related activity. (D) Proportion of non-responsive, movement-related, or reward-related neurons per FOV (n = 12 FOVs, N = 6 mice). Black dots represent individual FOVs, bars represent mean ± 95% CI. (E) Pairwise trial-to-trial correlation of population ΔF/F_0_ during push (blue) or pull (green) trials as a function of the pairwise trial-to-trial correlation of corresponding motion index (n = 12 FOVs, N = 6 mice). (F) Activity of two example movement-invariant neurons. *Top*, raster showing normalized ΔF/F_0_ across successive push (blue) or pull (green) trials; *Bottom*, mean ΔF/F_0_ ± 95% CI for push and pull trials. Dashed lines, movement initiation (MI). (G) Summary of movement-invariant and movement-bias neuron classification in layer 5B CFA (n = 468 neurons, N = 6 mice). (H) 2 overlapping FOVs from a single mouse showing movement-invariant (dark gray), movement-bias (light gray) and non-responsive neurons (white). (I) Activity of movement bias neurons split by type. *Top*, example Type 1 neurons with push (left) or pull (right) bias; *Bottom left*, example Type 2 neuron with push bias; *Bottom right*, example Type 3 neuron with pull bias. Dashed lines, movement initiation (MI). (J) Summary of movement bias classification in layer 5B CFA (n = 181 neurons, N = 6 mice). Insets, model examples of Type 1-4 ΔF/F_0_ changes. (K) 2 overlapping FOVs from a single mouse showing neurons with push (blue) or pull (green) bias. Gray, movement-invariant, non-responsive or reward phase neurons. (L) Mean pairwise trial-to-trial ΔF/F_0_ correlation for push (blue) and pull (green) trials in invariant, push- and pull-biased neurons (n = 468 neurons from 12 FOVs, N = 6 mice). Black dots represent individual mice. (M) Proportion of invariant, push- and pull-biased neurons per mouse (n = 468 neurons from 12 FOVs, N = 6 mice). Black dots represent individual mice. Red cross marks identified outlier.

### A small proportion of layer 5B neurons decode movement type

Next, we investigated how reliably movement-type could be decoded from single neuron and population changes in ΔF/F_0_ using a Gaussian Naïve Bayes classifier and logistic regression, respectively (see Methods). Approximately 37 % of neurons (172 / 468 neurons) displayed decoding accuracy scores above chance (Figure 3A), similar to, but slightly higher than the combined proportion of identified push and pull bias neurons (see Figure 2M), likely reflecting subtle differences in the sensitivity of both approaches. Given the trial-to-trial variability in ΔF/F_0_ and resultant moderate decoding scores (see Figure 2I and Figure 3A), we reasoned that population responses could provide a more robust movement-related signal that would enhance decoding of movement type. By applying logistic regression, population decoding was found to be consistently more accurate when compared to single neuron decoding (single cell median decoding accuracy = 0.61, [0.07] IQR; population median decoding accuracy 0.75, [0.16] IQR, N = 6 mice, *P* = 2.8×10^−2^, Wilcoxon signed rank test) (Figure 3B). However, this increase was driven almost entirely by a small proportion of high decoding accuracy neurons. Removing the top ∼20% of neurons ordered by decoding accuracy score abolished movement-type classification (median prop. removed = 0.21, [0.50] IQR, N = 6 mice) (Figure 3D-3E), whereas sequential removal of randomly selected neurons resulted in a significantly larger proportion of neurons having to be removed before decoding accuracy reduced to chance (median prop. removed = 0.64, [0.57] IQR, N = 6 mice, *P* = 2.8×10^−2^, Wilcoxon signed rank test) (Figure 3F). This dependency on high decoding accuracy neurons suggests that movement-specific information is routed through a selected subset of layer 5B neurons.

**Figure 3.**
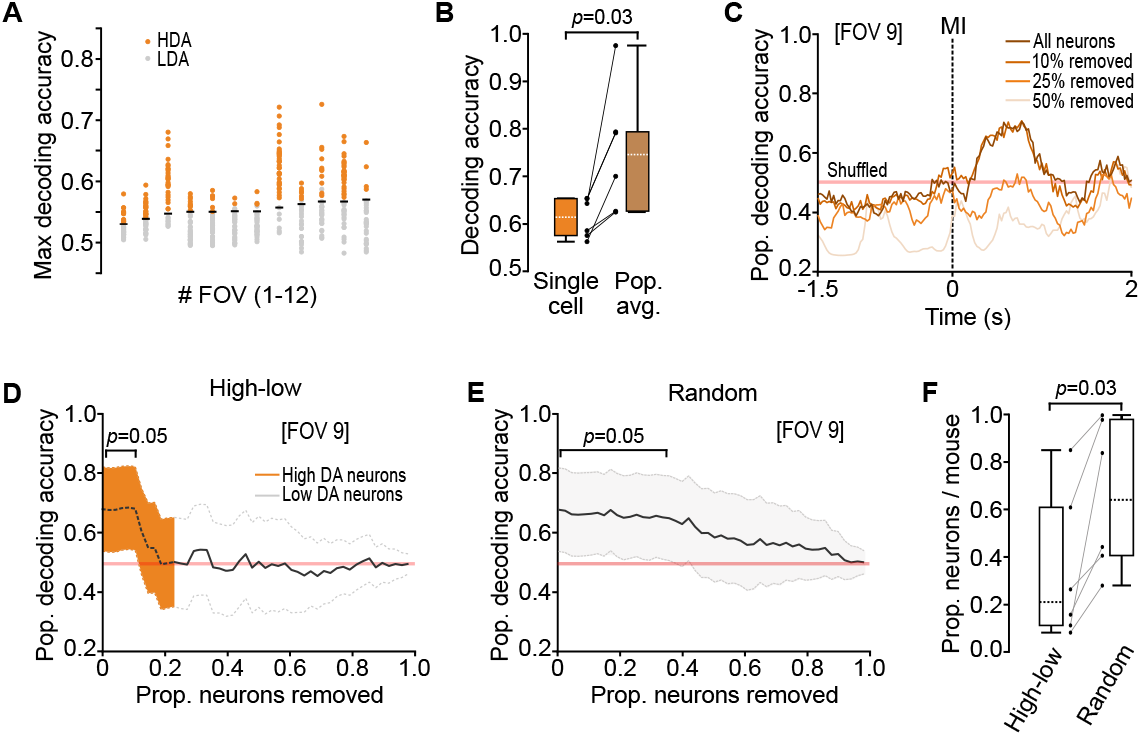
Population decoding relies on a small proportion of high decoding accuracy neurons. (A) Maximum decoding accuracy during peri-movement epochs generated using a Gaussian naïve Bayes classifier. Circles represent individual neurons; black horizontal lines indicate significance threshold. HDA, high decoding accuracy (orange); LDA, low decoding accuracy (gray). (B) Box-and-whisker plots showing median, interquartile range and range of single cell (naïve Bayes classifier, orange) and population (logistic regression, brown) decoding accuracy (n = 12 fields-of-view (FOVs), N = 6 mice, P = 2.8×10^−2^, Wilcoxon Signed-rank test). Black dots represent individual mice. (C) Mean population decoding accuracy for all neurons from a representative FOV or after removal of 10-50% neurons in order from high to low single cell decoding accuracy (Figure 3A). Red shaded line depicts the 95% CI based on shuffled data. Dashed line, movement initiation (MI). (D) Change in population decoding accuracy for an example FOV after the sequential removal of neurons in order from high (orange) to low (gray) single cell decoding accuracy (Figure 3A). Line, mean ± 95% CI. Red line, 95% CI based on shuffled data. (E) Change in population decoding accuracy for an example FOV after the random removal of individual neurons. Line, mean ± 95% CI. Red line, 95% CI based on shuffled data. (F) Box-and-whisker plots showing the median, interquartile range and range for ordered (high-to-low decoding accuracy) versus random removal of neurons (n = 12 FOVs, N = 6 mice, P = 2.8×10^−2^, Wilcoxon Signed-rank test). Black dots represent individual mice.

To further explore the underlying structure of layer 5B population activity, we employed principal component analysis (Churchland et al., 2012; Churchland et al., 2010; Cunningham and Yu, 2014; Kaufman et al., 2014; Stopfer et al., 2003). For the leading 16 principal components, we compared the difference between push and pull trials to compute a discrimination index (d’) (Figure S3A). Leading principal components tended to be more similar across actions, while action type was often better represented by higher components (Figure S3B). Despite correlating with population decoding scores, high d’ values were not preferentially associated with the leading principal components of the population activity (Figure S3D-3F), suggesting movement-type is not a dominant signal in the population response (Kaufman et al., 2016).

### IT and PT neurons display temporal differences in the encoding of movement type

Layer 5B contains two broad classes of projection neurons: intratelencephalic (IT) neurons form striatal and cortico-cortical connections (Levesque et al., 1996; Muñoz-Castañeda et al., 2021; Wilson, 1987; Winnubst et al., 2019); while pyramidal tract (PT) neurons target multiple subcortical, brainstem and spinal cord areas (Economo et al., 2018; Kita and Kita, 2012; Muñoz-Castañeda et al., 2021; Ueta et al., 2014; Winnubst et al., 2019) (Figure 4A). We next sought to understand whether movement-invariant and movement-specific signaling was dependent on projection class identity. To perform population imaging from identified cell types, we used an intersectional retrograde viral approach targeting ipsilateral brainstem (pons, PT) and contralateral CFA (IT) using a retrograde virus (r-AAV-retro cre) and conditional expression of GCaMP6s in ipsilateral CFA (Figure 4B-4C). Using a bicistronic viral vector expressing both GCaMP6s (flex) and mRuby, we confirmed that we recorded the majority of PT and IT neurons at depths up to 700 μm from the pial surface (Figure S4), consistent with our previous estimates (see Figure S2A-2C). Comparing push and pull trials, most PT neurons displayed movement-invariant activity (75.0 % [21.1] IQR), with a small number of neurons displaying push or pull bias (push bias = 10.3 % [19.9] IQR, pull bias = 13.8 % [11.3] IQR, N = 5 mice), mainly consisting of Type 1 (78.2 %) and Type 2 neurons (16.9 %) (Figure 4D-4E). In contrast, similar proportions of IT neurons displayed movement-invariant and movement specific signaling (movement-invariant = 51.2 % [11.6] IQR, movement-bias = 48.8 % [11.6] IQR), with Type 1 and Type 2 neurons again being the most abundant (Figure 4F-4G). Single cell decoding accuracy scores were highly consistent across mice and, as expected, population decoding accuracy increased when averaging the activity of all high decoding accuracy neurons per FOV (Figure 4H-4I). Although trial type could be decoded in approximately one third of projection neurons during the peri-movement window, the proportion of IT neurons with decoding accuracy above chance was highest prior to movement initiation, while for PT neurons this occurred during movement execution (IT peak proportion of neurons, 0.19 at -192 ms; PT peak proportion of neurons, 0.21 at +544 ms, N = 6 and 5 FOVs from 5 and 4 mice, respectively), suggesting differential roles for IT and PT neurons in movement initiation and execution, respectively. Importantly, at no time during the peri-movement window was the proportion of high decoding accuracy neurons above 21 % for either cell type (Figure 4J), consistent with a small proportion of projection neurons conveying time-dependent movement-specific information.

**Figure 4.**
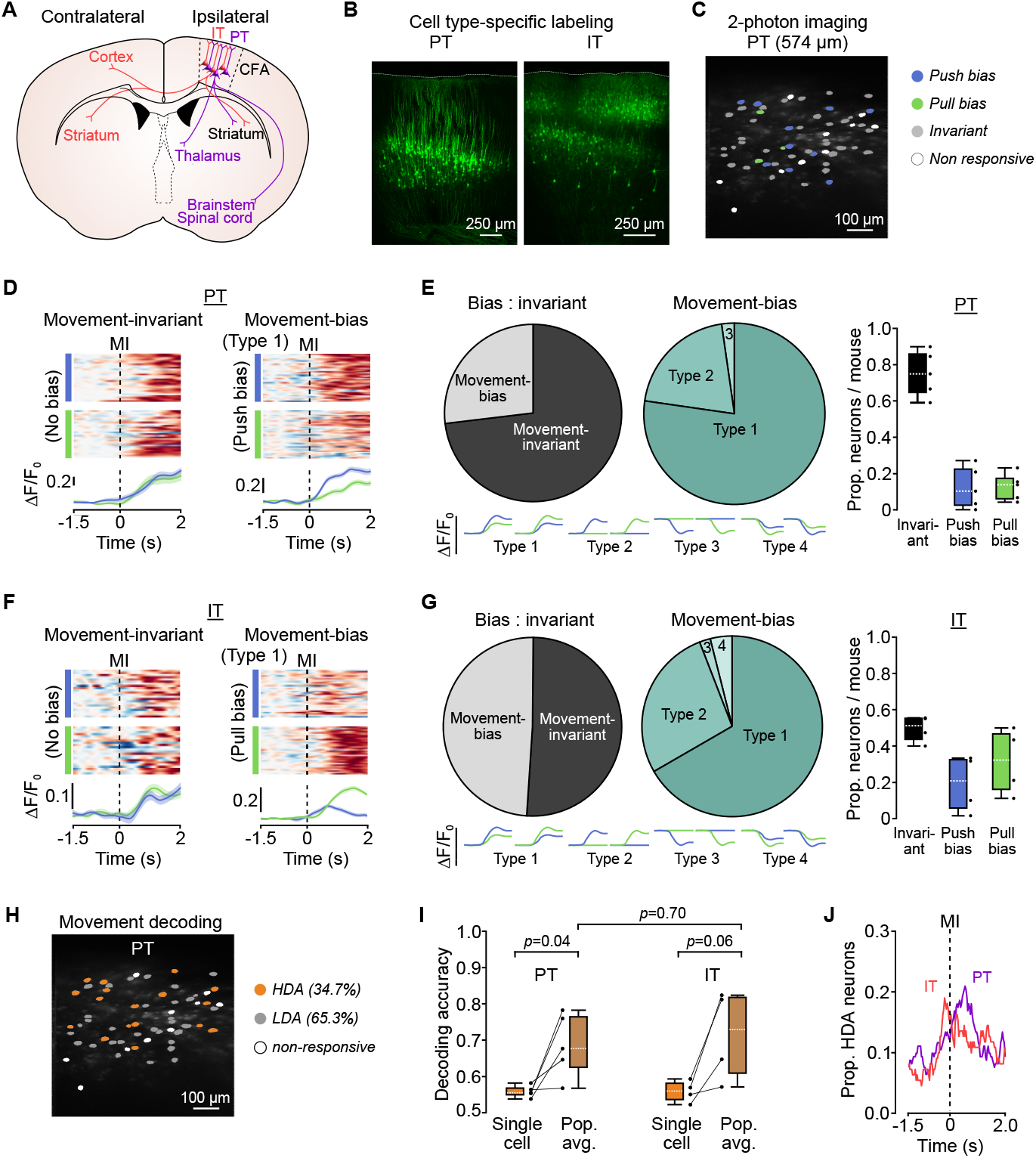
Movement-invariant and movement-specific signaling in identified layer 5B projection neurons. (A) Schematic showing brain-wide projections of layer 5B PT (purple) and IT (red) neurons. Contra- and ipsilateral relates to to the site of 2-photon imaging. (B) Histology from two imaged mice showing retrograde cell type-specific labeling of PT (left) and IT (right) neurons in CFA. (C) Example field-of-view (FOV) showing PT neurons with push (blue) or pull (green) bias. Gray, movement-invariant; white, non-responsive neurons. (D) Activity of two example PT neurons - movement-invariant (left) and movement-bias, Type 1 (right). *Top*, raster showing normalized ΔF/F_0_ across successive push (blue) or pull (green) trials; *Bottom*, mean ΔF/F_0_ ± 95% CI for push and pull trials. Dashed lines, movement initiation (MI). (E) *Left*, summary of movement-invariant and movement-bias PT neuron classification (n = 125 vs 46 neurons, N = 5 mice); *Middle*, summary of movement-bias classification in layer 5B CFA (n = 46 neurons, N = 5 mice). *Right*, proportion of invariant, push- and pull-biased neurons per mouse (n = 171 neurons from 6 FOVs, N = 5 mice). Black dots represent individual mice. *Below*, model examples of ΔF/F_0_ changes classified as Type 1-4. (F) Activity of two example IT neurons - movement-invariant (left) and movement-bias, Type 1 (right). *Top*, raster showing normalized ΔF/F_0_ across successive push (blue) or pull (green) trials; *Bottom*, mean ΔF/F_0_ ± 95% CI for push and pull trials. Dashed lines, movement initiation (MI). (G) *Left*, summary of movement-invariant and movement-bias IT neuron classification (n = 56 vs 54 neurons, N = 4 mice); *Middle*, summary of movement-bias classification in layer 5B CFA (n = 54 neurons, N = 5 mice). *Right*, proportion of invariant, push- and pull-biased neurons per mouse (n = 110 neurons from 5 FOVs, N = 4 mice). Black dots represent individual mice. *Below*, model examples of ΔF/F_0_ changes classified as Type 1-4. (H) Example FOV showing high (orange) and low (gray) decoding accuracy, and non-responsive (white) PT neurons. (I) Box-and-whisker plots showing median, interquartile range and range of single cell (naïve Bayes classifier, orange) and population (logistic regression, brown) decoding accuracy of PT (left) and IT (right) neurons. Comparisons made with two sample t-test. Black dots represent individual mice. (J) Proportion of neurons with decoding accuracy above chance (i.e., high decoding accuracy, HDA) across time. PT, purple; n = 58 / 171 neurons from 6 FOVs, N = 5 mice and IT, red; n = 43 / 110 neurons from 5 FOVs, N = 4 mice. Dashed line, movement initiation (MI).

### Movement-specific signaling is distributed across layer 5B

To explore whether high decoding accuracy PT and IT neurons form functional clusters within CFA, we first detected the onset of movement-related ΔF/F_0_ changes. Within each FOV, activity changes occurred ∼300 ms prior to movement, consistent with a role in preparation / initiation (Dacre et al., 2021; Estebanez et al., 2017; Isomura et al., 2009; Li et al., 2015), and tiled the peri-movement window up to reward delivery. Neurons displaying a range of ΔF/F_0_ onsets were spatially distributed across each FOV (Figure 5A-5B). To explore correlations in peri-movement activity patterns, we split PT and IT neurons based on their decoding accuracy scores (high, low and all) and compared pairwise activity during push and pull trials. We found weak correlations within and across groups irrespective of cell-type identity (Figure 5C-5D and Figure S5). Moreover, comparing the activities of PT and IT neurons as a function of their pairwise distance, suggested that neighboring neurons did not show correlated activity or spatiotemporal clustering (Figure 5E-5G). Thus, our data suggest a model whereby movement-specific information is routed through small, distributed subpopulations of layer 5B projection neurons, while most neurons convey movement-invariant information related to the timing of movement execution (Figure 5H).

**Figure 5.**
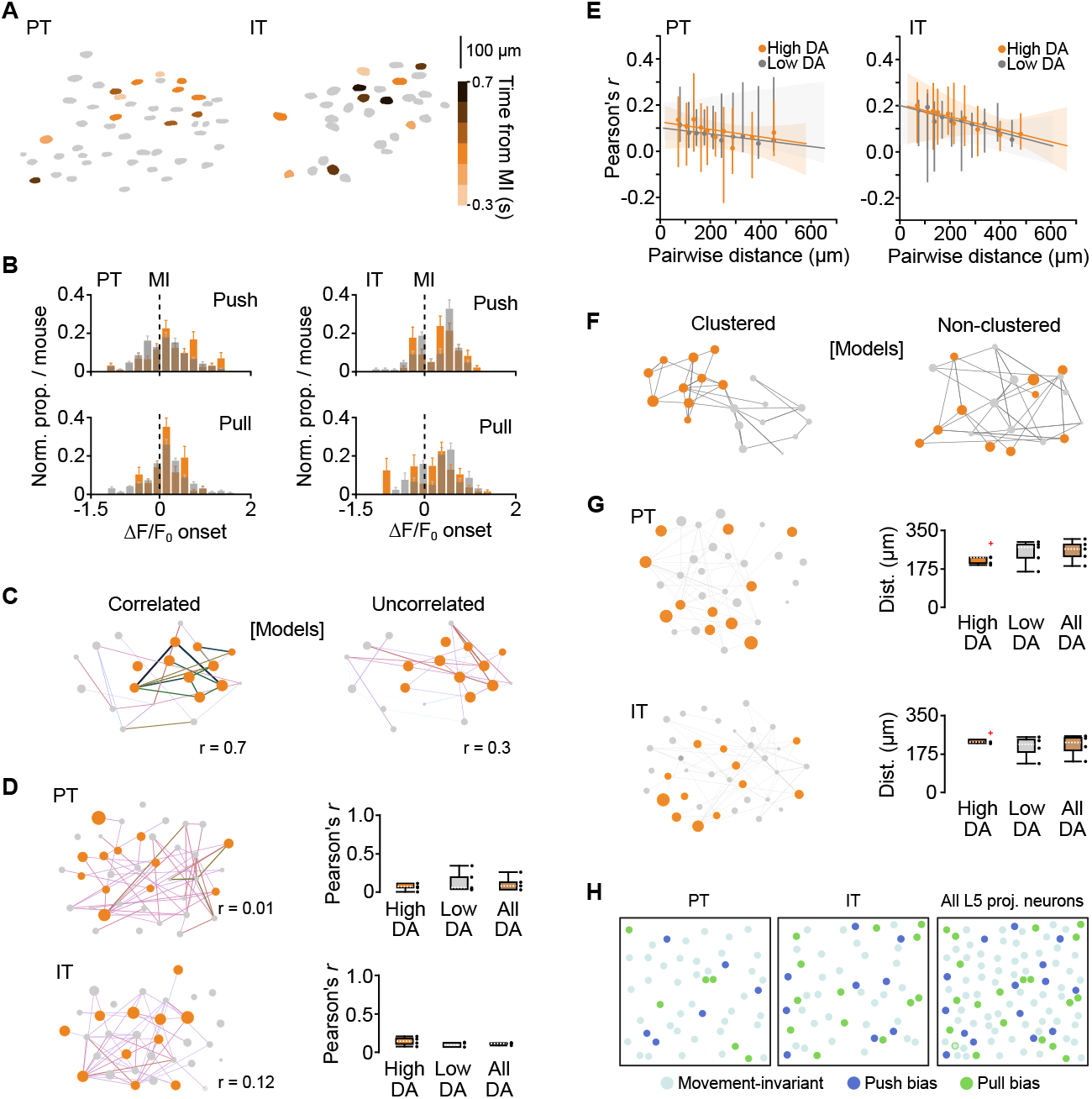
Cell type-specific spatiotemporal organization of high decoding accuracy neurons in layer 5B. (A) Example fields-of-view (FOVs) showing spatial distribution of ΔF/F_0_ onsets for high decoding accuracy PT (left) and IT (right) neurons during push trials. MI, movement initiation. Colors represent 200 ms bins tiling the peri-movement epoch: -300 ms (light orange) to +700 ms (dark brown). (B) Histograms of ΔF/F_0_ onsets for high decoding accuracy (orange) and low decoding accuracy (gray) PT (left) and IT (right) neurons during push (top) and pull (bottom) trials (n = 6 & 5 FOVs, N = 5 and 4 mice, respectively). (C) Modelled functional networks depicting high (orange) and low (gray) decoding accuracy neurons with correlated (left) or uncorrelated (right) activity. Each node, represented by a circle, corresponds to a neuron, while the connections represent the strength of activity correlation between neurons. (D) *Left top & bottom*, functional networks constructed from the pairwise activity correlations from a representative PT (top) and IT (bottom) FOV. Line color (light to dark) and width correspond to increasing values of Pearson’s *r*. Neurons are plotted as nodes in Euclidean space with color and size relating to increasing decoding accuracy. *Right top & bottom*, box-and-whisker plots showing the median, interquartile range and range of correlation coefficients across mice for high decoding accuracy (HDA, orange), low decoding accuracy (LDA, gray) and all (brown) PT (top) and IT (bottom) neurons. Black dots represent individual mice. (E) Median pairwise correlation coefficient with 95% CI as a function of pairwise distance for high decoding accuracy (orange) and low decoding accuracy (gray) PT (left) and IT (right) neurons. Horizontal lines denote linear regression model fit, with shaded regions representing the bootstrapped 95% CI (PT: P = 0.87 (HDA), P = 1.0 (LDA), n = 3024 observations, N = 5 mice; IT: P = 0.6 (HDA), P = 1.0 (LDA), n = 1562 observations, N = 4 mice). (F) Modelled functional networks depicting clustered (left) and non-clustered (right) high decoding accuracy neurons. Each node, represented by a circle, corresponds to a neuron, while the connections represent the pairwise distances between neurons. (G) *Left top & bottom*, functional networks constructed from the pairwise distances between neurons in a representative PT (top) and IT (bottom) FOV. *Right top & bottom*, box-and-whisker plots showing the median, interquartile range and range of median pairwise distances across mice for high decoding accuracy (HDA, orange), low decoding accuracy (LDA, gray) and all (brown) PT (top) and IT (bottom) neurons. Black dots represent pairwise distances for individual mice, red crosses mark identified outliers. (H) Models depicting cell type- and movement-specific layer 5B movement signaling in caudal forelimb motor cortex (CFA). Colored circles represent movement-invariant (cyan), push biased (blue) and pull biased (green) neurons.

## Discussion

Here, we show that the majority of layer 5B neuron signaling in mice was movement-invariant, where similar activity patterns were generated during the execution of two diametrically opposing movements. Changes in activity were tightly locked to the peri-movement period, indicative of a generic motor signal relating to movement timing, but not to movement type. Movement- or condition-invariant signaling also dominates in primate motor cortex, thought to trigger state-dependent switching from stable neural dynamics during rest towards oscillatory dynamics underpinning movement execution (Churchland et al., 2012; Churchland et al., 2010; Kaufman et al., 2014; Kaufman et al., 2016; Kurtzer et al., 2005) and is an emergent property of recurrent neural networks trained to recapitulate complex muscle patterns during reaching (Sussillo et al., 2015). In contrast to primate motor cortex, we found widespread movement-invariant responses at the single neuron level (Kaufman et al., 2016). This is unlikely to reflect differences in recording sensitivity, given that our imaging and electrophysiology approaches identified similar proportions of movement-invariant neurons across layer 5B (Wei et al., 2020; Zhou and Tin, 2021), or the limited number of movements in our task as movement-invariant responses have been shown in relatively simple tasks requiring few actions (Evarts, 1968; Hocherman and Wise, 1991; Messier and Kalaska, 2000; Riehle et al., 1994; Weinrich et al., 1984). Instead, this might reflect evolutionary differences in how motor cortex recruits and controls muscle activation during the transition from rest to movement execution. Cell-type-specific imaging identified that movement-invariant signaling dominated PT neuron activity, suggesting a large proportion of the output conveyed to subcortical, brainstem and spinal cord areas relates to the execution of movement without necessarily specifying movement type, whereas equal proportions of IT neurons displayed movement-invariant versus movement-specific signaling. If movement-invariant signaling relates to the execution of movement and dominates deep layer motor cortex activity, what drives the change in neural activity and what purpose might it serve? Long-range inputs from basal ganglia, secondary motor cortex and cerebellum are possible sources (Hooks et al., 2013; Hooks et al., 2018; Nelson et al., 2021), providing an external trigger to transform motor cortical dynamics necessary for postural maintenance at rest to a neural state required for movement execution. This switch in neural dynamics would signify the intention to move, but not which movement will be executed (Elsayed et al., 2016; Kaufman et al., 2016). An important next step will be to develop methods to identify and selectively manipulate neurons displaying movement-invariant signaling to demonstrate their causal contribution to postural control and timing of movement initiation.

We reasoned that the execution of two diametrically opposing movements should, in principle, generate distinct patterns of cortical output dynamics, given differences in starting posture, direction of movement and temporal sequence of muscle activation (Isomura et al., 2009; Miri et al., 2017). Although we found that the majority of layer 5B neuron signaling was movement-invariant, a relatively small proportion of neurons displayed response bias towards either push or pull movements. The low prevalence of movement-specific signaling is unlikely to be due to masking of subtle changes in spike rate when using calcium reporters (Wei et al., 2020; Zhou and Tin, 2021), since we observed similar proportions of movement-specific signaling when performing high-density extracellular recordings of putative layer 5B projection neurons. The firing rates of individual neurons in motor cortex reflect a complex combination of signals that have been shown to correlate with joint angle, direction, and speed (Georgopoulos et al., 1982; Moran and Schwartz, 1999; Paninski et al., 2004; Thach, 1978), while population dynamics reflect time-varying changes in neural state during the transition from rest to movement execution (Churchland et al., 2012; Churchland et al., 2010; Kaufman et al., 2014; Kaufman et al., 2016; Kurtzer et al., 2005; Sauerbrei et al., 2019). In mice, individual layer 5B neurons displayed moderate decoding accuracy scores, likely due to relatively high trial-to-trial variability, while the population average was consistently higher across mice. We found that only a small proportion (∼20%) of neurons contributed to high population decoding accuracy scores, with their combined effects accurately decoding three quarters of all trials. Removing only a handful of neurons per FOV was sufficient to abolish decoding, confirming that a minority of neurons convey the majority of information regarding movement type. This dependency on only a few neurons has important implications for understanding how movement-specific information is encoded in primary motor cortex, given that mapping neural dynamics during the execution of a single movement task will likely uncover widespread movement-invariant signaling, that relates to limb movement, but not the specific movement being executed.

In mouse cortex, projection neurons display connectivity patterns both within and across classes that suggest general organizing principles (Brown and Hestrin, 2009; Kiritani et al., 2012; Maruoka et al., 2017; Morishima et al., 2011). IT neurons in motor cortex are strongly recurrently connected, whereas inter-class connectivity is largely directional from IT to PT but not vice versa, generating a hierarchical organization with unidirectional signaling from higher-order to lower-order output neurons (Kiritani et al., 2012). Asymmetric projection-class connectivity, as well as differences in input structure and intrinsic excitability (Anderson et al., 2010; Hooks et al., 2013; Kiritani et al., 2012; Oswald et al., 2013) provide a mechanism to flexibly route movement-specific information via two independent output channels depending on behavioral requirements. Our cell-type-specific imaging identified that only a small proportion of PT neurons conveyed movement-specific information. In contrast, almost half of IT neurons displayed movement-specificity, with similar proportions of both push and pull bias. Although PT and IT activity onsets occurred prior to and throughout movement, consistent with both pathways contributing to movement initiation and execution (Chen et al., 2017; Economo et al., 2018; Li et al., 2015; Park et al., 2021; Wang et al., 2017), the proportion of IT neurons with high decoding accuracy was highest prior to movement initiation, while for PT neurons this occurred during movement execution. This suggest that information relating to movement type is first conveyed by IT neurons, which project to bilateral striatum and cortex, but not other subcortical targets (Levesque et al., 1996; Muñoz-Castañeda et al., 2021; Wilson, 1987; Winnubst et al., 2019), before PT neurons then propagate information to subcortical, brainstem and spinal cord circuits necessary for online control of forelimb movement (Economo et al., 2018; Kita and Kita, 2012; Muñoz-Castañeda et al., 2021; Ueta et al., 2014; Winnubst et al., 2019). Importantly, the proportions of PT or IT neurons decoding movement type never exceeded 25%, consistent with movement-specific signaling being confined to a relatively small subpopulation of layer 5B projection neurons. What is unclear is the extent to which movement-specific signaling in PT and IT neurons is organized by projection target as seen in anterolateral motor cortex during directional licking (Chen et al., 2017; Economo et al., 2018; Li et al., 2015). Targeting neurons based on molecular expression profiles and projection specificity (Muñoz-Castañeda et al., 2021; Winnubst et al., 2019) will provide a finer grained understanding of how movement-specific information is routed via molecularly distinct projection pathways.

We found that PT and IT neurons displaying high decoding accuracy were distributed across FOVs. This lack of functional clustering differs from the proposed modular organization of directionally tuned cells in primate motor cortex, where neurons with similar preferred direction tend to cluster into vertically oriented minicolumns approximately 50 - 100 μm wide, repeated every 250 μm (Amirikian and Georgopoulos, 2003; Cheney et al., 1985; Georgopoulos et al., 2007; Jones and Wise, 1977), but consistent with the distributed spatial organization of direction-specific layer 5B projection neurons in mouse anterolateral motor cortex during the execution of a whisker-based object location discrimination task (Li et al., 2015). The apparent lack of spatial clustering in CFA is unlikely to be due to reduced sensitivity of our analysis methods as confidence intervals provide a lower bound indication of cluster size such that, if present, spatial clusters based on decoding accuracy would have to be less than ∼50 μm. Similarly, we found no evidence of temporal clustering in either high (movement-specific) or low (movement-invariant) decoding accuracy neurons, as expected given that the onsets of both PT and IT neuron activity changes occurred ∼300 ms prior to movement and tiled the peri-movement period up to reward delivery. Our work extends previous findings in superficial layers of motor cortex showing neurons with task-related response properties are spatially intermingled (Galiñanes et al., 2018; Huber et al., 2012), to support a model in which movement-specific signaling in layer 5B is routed via small but distinct subpopulations of projection neurons. The flexible routing of information through distributed descending projection pathways could in principle provide a mechanism for differentially controlling movement variables necessary for the execution of a large repertoire of limb movements.

## Supporting information

Video - example cued push and pull trials

Video - muscimol inactivation of CFA

## Acknowledgments

We are grateful to Gülsen Sürmeli, Matt Nolan and members of the Nolan, Duguid and Sürmeli labs for experimental discussions and for comments on the manuscript. Nick Steinmetz for the suggested design of the author contribution matrix. AAV-GCaMP6s was a gift from Douglas Kim & GENIE Project (Addgene 100844-AAV1). AAV-pgk-Cre was a gift from Patrick Aebischer (Addgene viral prep # 24593-AAVrg). pAAV-CAG-Flex-mRuby2-GSG-P2A-GCaMP6s-WPRE-pA was a gift from Tobias Bonhoeffer & Mark Huebener & Tobias Rose (Addgene viral prep # 68717-AAV1). Confocal microscopy was performed in the IMPACT Imaging Facility at the University of Edinburgh. Research was supported by grants from the Biotechnology and Biological Sciences Research Council (BB/R018537/1 to I.D.), DFG fellowship program (AM 443/1-1 to J.A.), the Shirley Foundation, Simons Initiative for the Developing Brain (SIDB) PhD studentship (C.E.), A*STAR PhD studentship (B.P.) and a Wellcome Senior Research Fellowship (110131/Z/15/Z to I.D.).

## Author contributions

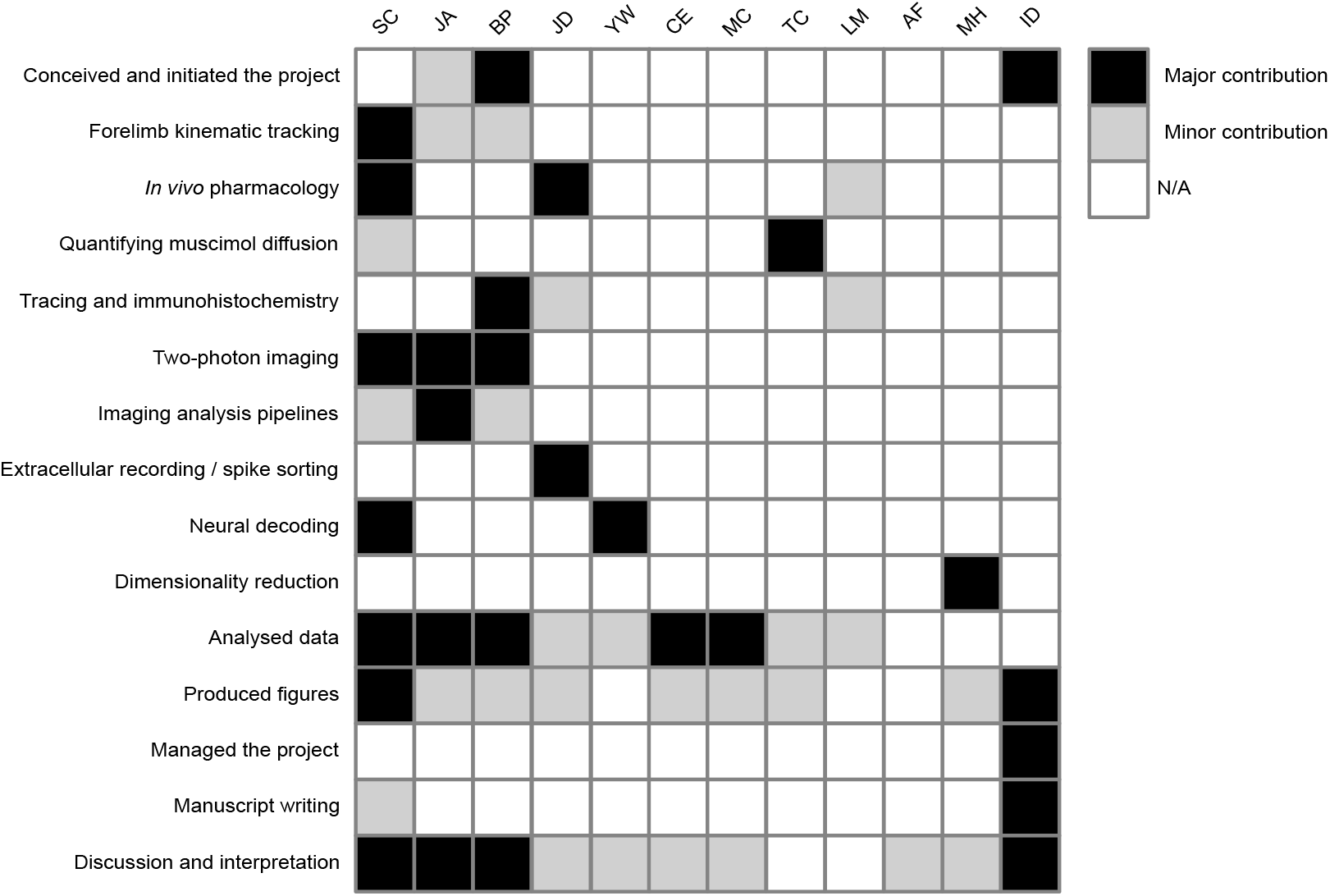

## Methods

### Animal husbandry and general surgery

Male adult C57BL/6J wild-type mice (5-12 weeks old, 20-30 g, 1-4 animals per cage) were maintained on a reversed 12:12 hour light:dark cycle and provided ad libitum access to food and water as well as environmental enrichment. All experiments and procedures were approved by the University of Edinburgh local ethical review committee and performed under license from the UK Home Office in accordance with the Animal (Scientific Procedures) Act 1986. Surgical procedures were performed under ∼1.5% isoflurane anaesthesia and each animal received fluid replacement therapy (0.5ml sterile Ringer’s solution), buprenorphine (0.05 mg/kg) and either carprofen (4 mg/kg) or dexamethasone (2 mg/kg) for pain relief and to reduce inflammation. At 24 and 48 hours, carprofen (4 mg/kg) was administered for post-operative pain relief. Craniotomies were performed in a stereotactic frame (David Kopf Instruments, CA, USA) using a hand-held dentist drill with 0.5 mm burr. A small lightweight headplate (0.5 g) was implanted on the surface of the skull using cyanoacrylate glue and dental cement (Lang Dental, IL, USA) and mice were left for at least 48 hours to recover.

### Behavioral training

After recovery from head plate surgery, mice were handled extensively before being head restrained and habituated to a custom forelimb lever push / pull behavioral setup. Mice were trained to perform two diametrically opposing movements - a 4 mm push or pull of a moveable lever - in response to a 6 kHz auditory cue to obtain a 5 μl water reward. Mice rested their right forepaw on a stationary lever while making push or pull movements with their left forepaw. Upon completion of a successful push or pull (determined by the status of IR beams at either end of the lever’s travel), the moveable lever was locked in place for the duration of the reward period (3 s) and the water reward was delivered by an automated spout - both locking mechanism and spout were controlled by micro servo motors (HXT900, HexTronik). To increase task engagement, mice were placed on a water control paradigm (1 ml/day) and weighed daily to ensure body weight remained above 85% of baseline. Mice were trained once per day for 30 mins and advanced through different phases of the task once they achieved > 50 rewards in each of two consecutive sessions or > 70 rewards in a single session. Initially, mice were required to perform uncued push and pull movements to obtain rewards (phase 1). Next, an auditory cue was introduced with pseudo-random inter-trial-interval (ITI) of 4-6 s and a response window of 10 s (phase 2). During the ITI, mice had to hold the moveable lever still - spontaneous movements of the lever within the ITI triggered a reset, where the lever was locked in the original position for 1 s. The response window was gradually reduced to 5 (phase 3) and then 2 s (phase 4) across training sessions. Mice were deemed “expert” after achieving > 70 rewards on two consecutive days of training with a response window of 2 s. During 2-photon imaging experiments (see below), a 1 s delay between completion of a successful movement and reward delivery was introduced to temporally separate activity related to reward consumption from the forelimb movements of primary interest.

### Forelimb kinematic tracking

Behavior was recorded using a high speed camera (60 fps during population calcium imaging of layer 2/3 or 5B - Prosilica GC780C, Allied Vision, Germany; or either 100 or 300 fps for cell-type specific calcium imaging or in vivo pharmacology, respectively - Mako U U-029, Allied Vision) and acquired using Streampix 7 (Norpix, Canada) or Mantis64 (https://www.mantis64.com/). To measure gross forelimb movement, a region of interest (ROI) was manually drawn around the left forelimb and the frame-to-frame difference in pixel intensity was calculated using the formula: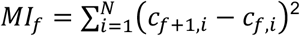, where c_f,i_ is the grayscale level of pixel i in frame f. The resulting motion index was smoothed with a 1 s LOESS filter then aligned to behaviorally relevant time points (lever displacement, cue presentation etc.), with videos and behavior resampled to a common sampling rate. Motion index onsets were calculated by aligning the motion index to the lever movement and identifying the first point prior to movement where mean motion index was > threshold (mean upper bound of 95% confidence interval during baseline). Directional tracking of forelimb and lever movement was performed using Deep Lab Cut, a markerless video tracking toolbox (Mathis et al., 2018). Tracking data was aligned to cue presentation, baselined to mean xy position during the 100 ms prior to cue and then cropped between movement initiation and movement completion. For presentation, trials of different durations were resampled to a fixed length to enable a mean trajectory to be plotted across multiple trials.

### In vivo pharmacology

To assess the behavioral effects of caudal forelimb area (CFA; N = 10) or hind limb motor cortex (M1_hl_; N = 5) inactivation, mice trained to expert level in the task had a small burr hole opened directly above the target area (CFA: 1.6 mm lateral, 0.6 mm rostral to bregma; M1_hl_: 1.25 mm lateral, 1.25 caudal to bregma) before recovering for > 90 mins. After 10 mins of baseline behavioral task execution, the lever was locked and a small volume of the GABA_A_ receptor agonist muscimol (dissolved in external saline solution containing 150 mM NaCl, 2.5 mM KCl, 10 mM HEPES, 1.5 mM CaCl_2_ and 1 mM MgCl_2_, adjusted to pH 7.3) was injected into the target area - 200 nl of 2mM muscimol was injected at each of 4 or 5 sites forming a 0.5 mm X centered on 1.6 mm lateral, 0.6 mm rostral to bregma (CFA) or 1.25 mm lateral, 1.25 caudal to bregma (M1_hl_). Each injection site was at a depth of 0.7 mm below the cortical surface. To confirm the anatomical location of drug injection, 1% w/v of retrobeads (red, Lumaflor Inc.) was included in the injected solution. In a subset of mice (N = 5 of 10), animals were also injected in CFA with saline (vehicle only; injection performed on a different day). In these experiments, mice were randomly assigned to drug or control groups (each mouse received one injection of muscimol and one injection of saline) and experiments performed blinded. After each experiment, mice were transcardially perfused and coronal (60 μm) or sagittal (100 μm) sections were cut with a vibratome (Leica VT1000S), mounted with Vectashield mounting medium (H-1000, Vector Laboratories), imaged using a fluorescence microscope (Leica DMR, 5x objective) and manually referenced to the Paxinos and Franklin Mouse Brain Atlas (Paxinos and Franklin, 2008). Behavioral metrics were analyzed comparing videos of 10 mins pre and post injection - the experimenter was blinded to the experimental groupings. Behavioral video data for all pharmacology experiments was captured using a high-speed camera (Mako U U-029, Allied Vision), and paw position accuracy was calculated as the proportion of trials in which mice were holding the moveable lever with their left forepaw at the onset of the auditory cue.

### Quantifying muscimol diffusion

To measure muscimol diffusion, a small volume of muscimol-BODIPY TMR-X Conjugate (ThermoFisher Scientific; dissolved in 0.1PBS w/1% dimethyl sulfoxide) was injected into CFA (200 nl of 2 mM at 4 sites centered on 1.6 mm lateral, 0.6 mm rostral to bregma at a depth of 0.7 mm below the cortical surface). To mark the center of injection, pipettes were backfilled with a small volume (∼20 nl) of green retrobeads (Lumafluor Inc.) prior to filling with muscimol-BODIPY. Following injection, animals were transcardially perfused and brains snap-frozen on dry ice 10 minutes from completion of the muscimol injection. Brains were stored on dry ice, coronal sections (60 μm) collected with a cryostat (Leica) at -20C and imaged with a light microscope (Leica DMR, 5x objective). We assumed maximum fluorescence ≈ maximum injected concentration and that grayscale pixel intensity was proportional to muscimol-BODIPY concentration. Therefore, pixel values were thresholded at the equivalent pixel value of an EC_20_ concentration of muscimol and fluorescence outlines were drawn to generate a ‘spread profile’. Green retrobeads were used to mark the center of each injection, and images were aligned to the injection center of gravity. From the aligned profiles, a modal spread profile (i.e., pixels with positive grayscale values across all mice) was generated and aligned to the Paxinos and Franklin Mouse Brain Atlas (Paxinos and Franklin, 2008).

### Retrograde tracing

To selectively label pyramidal tract (PT) neurons in layer 5B of CFA, red retrobeads (Lumafluor, USA) were injected into the pons (4.0 mm caudal and 0.4 mm lateral to bregma ipsilateral to the target CFA), delivered via pulled glass pipettes (5μl, Drummond Scientific, PA, USA; 10-20 nl/min) using an automated injection system (Model Picospritzer iii, Parker, NH, USA). Injections were made at 4 sites (100 nl / site) located 200, 400, 600 and 800 μm dorsal from the cranial floor. After > 14 days post-injection, mice were terminally anaesthetized using an intraperitoneal injection of a ketamine/domitor mixture (75 mg/kg ketamine, 1 mg/kg domitor) and transcardially perfused with 30 ml of phosphate-buffered saline (PBS) followed by 30 ml of 4% paraformaldehyde (PFA, Sigma-Aldrich, MO, USA), dissolved in PBS and adjusted to pH 7.4. Brains were post-fixed at 4 °C for 1-3 days in 4% PFA solution, then transferred to PBS solution. Individual brains were cut into coronal sections (60 μm) using a vibrating microtome (Leica VT1200 S, Leica Microsystems (UK) Ltd.) and mounted with Vectashield Antifade Mounting Medium (Vector Laboratories, CA, USA). Images were acquired with a Leica DM R epifluorescence microscope and image analysis was performed using ImageJ (Rueden et al., 2017) and MATLAB (MathWorks, MA, USA). To obtain estimates of the depth of layer 5B in CFA, 3 coronal sections from each brain were imaged (0.54 mm, 0.6 mm and 0.66 mm rostral to bregma). Brightness/contrast adjustments and background subtraction (rolling ball, 30 pixels wide at 1.28 μm/pixel; Fiji (Schindelin et al., 2012)) functions were performed to reduce the contribution of background autofluorescence. Each ROI was then divided into 25 μm deep bins that were normalized to a value between 0 and 1, with 0 being the darkest bin and 1 being the brightest bin and all bins were compared to baseline. In order to obtain a depth profile of layer 5B within CFA, the depth of the dorsal-most retrogradely labeled neuron was recorded at 100 μm intervals from 1.3 - 1.9 mm lateral to bregma and repeated in 5 sequential coronal sections from 0.36 - 0.84 mm rostral to bregma. For each mouse, the depth of layer 5B at the center of CFA (1.6 mm lateral, 0.6 mm rostral to bregma) was taken as the reference depth and the depths of other locations reported relative to this value.

### Immunohistochemistry

Mice expressing GCaMP6s were transcardially perfused and horizontal sections (30 μm) were cut parallel to the surface of CFA. Sections were rinsed in PBS overnight, incubated with a blocking solution (PBS, with 0.5% Triton X-100 (Sigma-Aldrich) and 10% goat serum (Sigma-Aldrich)) for 2 hrs and rinsed with PBS. Sections were incubated overnight with mouse anti-NeuN (MAB377 Anti-NeuN Antibody, clone A60, Sigma-Aldrich) diluted 1:1500 in carrier solution (PBS, with 0.5% Triton X-100 and 1% goat serum), then rinsed with PBS. For secondary antibody binding, sections were incubated overnight with goat anti-mouse Alexa Fluor 568 (Invitrogen, MA, USA) diluted 1:750 in carrier solution then rinsed with PBS. Sections were mounted onto glass slides, briefly air-dried, covered with Vectashield Antifade Mounting Medium (Vector Laboratories), and sealed with a glass coverslip. Images of CFA were acquired using a Nikon A1R FLIM confocal microscope (20X objective lens, 0.8 NA, Plan Apo VC, Nikon, Europe). Three images were taken at imaging planes corresponding to layer 5B (550 μm from the cortical surface). The number of cells in each image was manually counted and divided by the area to obtain a measure of neuron density. For most fields-of-view (FOVs) recorded during calcium imaging, neurons were not visible in all portions of the frame due to occlusion by blood vessels. Polygons were manually drawn around visible neurons in each field-of-view to provide a realistic estimate of the imaging area.

### 2-photon imaging

To perform population calcium imaging in layer 5B (12 FOVs, N = 6 mice) of CFA, 200 nl of the adeno-associated virus (AAV) AAV1-SynGCaMP6s (diluted to 2.9×10^12^ GC/ml, Addgene 100844-AAV1) was injected into right motor cortex (1.6 mm lateral, 0.6 mm rostral to bregma and 0.6 mm from the cortical surface) via pulled glass pipettes (5 μl, Drummond Scientific; 10-20 nl/min) using an automated injection system (Model Picospritzer iii, Parker), before sealing the craniotomy with silicone (Body Double; Smooth-On, PA, USA) and implanting a lightweight headplate. For imaging, a cranial window (a glass coverslip - #0; Menzel-Gläser, Germany - fixed in place with cyanoacrylate glue), was implanted above the virus injection site. 2-photon calcium imaging was performed using a custom-built resonant scanning 2-photon microscope (320 × 320 μm field-of-view; 600 × 600 pixels) with a Ti:Sapphire pulsed laser (Chameleon Vision-S, Coherent, CA, USA; < 75 fs pulse width, 80 MHz repetition rate) tuned to 920 nm wavelength. Images were acquired at 40 Hz with a 40x objective lens (0.8 NA; Nikon) with custom-programmed LabVIEW-based software (LotoScan).

For cell type specific imaging, AAV-pkg-Cre (Addgene 24593-AAVrg; 1.7×10^13^ GC/ml) was injected into either the ipsilateral (right) pons (PT; 0.4 mm lateral, 0.4 mm rostral to lambda and 0.2, 0.4 and 0.6 mm dorsal from the cranial floor) or contralateral (left) CFA (IT; 4 injections centered on 1.6 mm lateral, 0.6 mm rostral relative to bregma and at two depths - 0.7 and 0.35 mm from the cortical surface) followed by an injection of pAAV-CAG-Flex-mRuby2-GSG-P2A-GCaMP6s-WPRE-pA (Addgene 68717-AAV1; 1.8×10^13^ GC/ml) into right motor cortex (1.6 mm lateral, 0.6 mm rostral to bregma and 0.6 mm from the cortical surface). 2-photon calcium imaging was performed using an 8 kHz resonant scanning microscope (HyperScope, Scientifica, UK; 690 × 690 μm field-of-view; 512 × 512 pixels) with a Ti:Sapphire pulsed laser (Chameleon Vision-S, Coherent, CA, USA; < 75 fs pulse width, 80 MHz repetition rate) tuned to 920 nm wavelength. Images were acquired at ∼30 Hz with a 16x objective lens (0.8 NA; Nikon) with SciScan image software (Scientifica) and synchronized with external high speed videos and behavioral data using Mantis64. To facilitate reliable imaging at depths > 500 μm, all imaging was performed acutely following cranial window implantation - usually within 24 hours of the surgery. Laser power was 91 – 173 mW (mean = 143 mW) across all imaging sessions, well below the damage thresholds of 250 – 300 mW outlined in Podgorski and Ranganathan (2016). The combination of low pixel dwell time and systematic blanking of field-of-view edges, where the dwell time is higher, and the addition of room temperature artificial cerebrospinal fluid on the surface of the skull further reduces the risk of thermal effects (as discussed in Podgorski and Ranganathan 2016).

Motion artifacts in the raw fluorescence videos were corrected using discrete Fourier 2-dimensional-based image alignment (SIMA 1.3.2; (Kaifosh et al., 2014)). ROIs were drawn manually in Fiji and pixel intensity within each ROI averaged to produce a raw fluorescence time series (F). To remove fluorescence originating from neuropil and / or neighboring neurons, fluorescence signals were decontaminated and extracted using nonnegative matrix factorization, as implemented in FISSA (Keemink et al., 2018). Normalized fluorescence was calculated as ΔF/F_0_, where F_0_ was calculated as the 5^th^ percentile of the 1 Hz low-pass filtered raw fluorescence signal and ΔF = F-F_0_. All further analyses were performed in custom written scripts in MATLAB or Python 3 that are available via the Duguid Lab GitHub repository (https://github.com/DuguidLab). Prior to further analysis, data were smoothed with a 1 s LOESS filter, chosen as it reliably captured the overall shape and peak response of the ΔF/F_0_ time series. In order to define movement-related neurons, we first defined baseline (−1.5 to - 0.5 s relative to motion index onset) and peri-movement (−0.15 s from motion index onset to +1.5 s after movement completion to capture all neuronal responses up to and including reward delivery) epochs. Movement-related neurons were identified by two independent methods: 1) a bootstrapped distribution (10,000 samples) of baseline-to-peak values (mean of the 100 ms centered on the largest deviation from baseline within the peri-movement epoch - mean of baseline epoch) was used to test whether 95% confidence intervals for the responses were different from 0; 2) bootstrapped distributions of mean ΔF/F_0_ in 250 ms bins within the peri-movement epoch were compared to bootstrapped distributions of mean ΔF/F_0_ within the baseline epoch. If either method identified significant differences the neuron was classified as movement-related. Neurons with no differences between baseline and movement epochs were classified as non-responsive and excluded from further analysis. Movement-related neurons with a median onset occurring after median movement completion (plus a small correction factor of 40 ms, to account for the rise time of GCaMP6s) were defined as ‘reward phase’ neurons and also excluded from further analysis. The median onset time of each cell was calculated by employing a previously published onset detection algorithm using a slope sum function (SSF;Zong et al., 2003; Dacre et al. 2021) with the decision rule and window of the SSF adapted to the calcium imaging data (threshold 10% of peak, SSF window 375 ms, smoothed with a Savitzky Golay filter across 27 frames with order 2) and reported as the median of 10,000 bootstrapped samples to reduce the influence of noisy individual trials. Prior to extracting ΔF/F_0_ onsets, we verified this algorithm with simulated data thereby accounting for any bias in the onset detection potentially introduced by filtering and/or the decision rule. To simulate the rising phase of the movement related calcium events in our data we used linear ramps with defined onset times and a rise time of 0.5 s mimicking GCaMP6s kinetics. We then calibrated the onset detection algorithm on the simulated data (100 simulated cells with 30 simulated trials per cell and artificially added noise in each trial matching the noise level in the imaging data) and updated it by a small correction factor. Neurons with movement bias were detected using the same classification criteria described above but across movements (i.e. significant differences in bootstrapped ΔF/F_0_ baseline-to-peak or 250ms peri-movement bins). Where significant differences between push and pull trials were identified, the neuron was classified as biased for the movement with the larger response from baseline.

### Trial to trial correlations

To assess the similarity of trial-to-trial activity, the average pairwise trial-to-trial correlation coefficients (Pearson’s r) of the peri-movement ΔF/F_0_, smoothed with a 1 s LOESS filter, were computed for each neuron. Data are presented as bootstrapped medians per animal for each movement preference class (10,000 repetitions, 50 samples). To investigate the relationship between trial-to-trial similarity of movement and population ΔF/F_0_, pairwise trial-to-trial correlation coefficients (Pearson’s r) of peri-movement motion index and pairwise trial-to-trial correlation coefficients (Pearson’s r) of the peri-movement population ΔF/F_0_ of the same trials were compared. Population ΔF/F_0_ was the sum of all movement-related neurons in each FOV. Data were binned according to the pairwise trial-to-trial correlation coefficients of their motion index and are presented as bootstrapped median (10,000 repetitions, 50 samples) within each bin.

### Extracellular recording and spike sorting

To compare neural activity during the task, extracellular unit recordings in CFA were performed using acutely implanted silicone probes (Neuropixels Phase 3B probes, IMEC). Data were acquired from the 384 channels closest to the probe tip (bank 0) with SpikeGLX software at 30 KHz, an amplifier gain of 500 for each channel and high-pass filtered with a cutoff frequency of 300 Hz. Spike data were synchronized with external high-speed videos and behavioral data (cue presentation, lever movement and reward delivery) using Mantis64. Spike sorting was performed using Kilosort3 to automatically cluster units from raw data (Pachitariu et al., 2016). The resulting spike clusters were manually curated using Phy (https://github.com/cortex-lab/phy), and any unit with sufficient refractory period violations, inconsistent waveform amplitude across the duration of the recording, or clipped amplitude distribution was excluded from analyses. Probe location was confirmed via DiI (Thermofisher) reconstruction of the recording track and compared to retrogradely labeled PT neurons - FastBlue (Polysciences) injected into the pons - in each animal to limit analysis to units within layer 5B (upper boundary, 500-680 μm; lower boundary, 900-1080 μm; N = 5 mice). Spike widths were calculated as the duration from trough to following maxima of the spike waveform. Putative pyramidal neurons were identified as units with median spike widths greater than 0.4 ms.

To classify units as responsive to the push or pull movements, firing rates were calculated by convolving motion index-aligned spike times with a 50 ms Gaussian kernel and mean changes in firing rate were calculated by subtracting the mean firing rate during a baseline period (1 s period before cue presentation) from the mean firing rates in 250 ms bins tiling a response period extending back from *max* (_*X*_*pushcompletion*, _*X*_*pullcompletion*)) to include motion index onset. Briefly, motion index onsets were calculated as the first point after cue where the motion index was > threshold (threshold = mean motion index in a baseline window from the 1.5 s before cue plus 2 SD). Trials where the motion index onset came before cue presentation were excluded from analysis. Significant responses were identified by comparing bootstrapped 95% confidence intervals of these mean changes in firing rates to 0; if at least one bin differed from 0, that unit is considered movement-responsive. Movement-responsive units were classified as having a push or pull bias if the confidence intervals in the earliest significantly responsive bin did not overlap; in these cases the units were classified as biased for the movement with the larger change from baseline.

### Neural decoding

To decode movement type in single neurons we employed a naïve Bayes classifier, where distributions of features are assumed to be Gaussian. Movement-aligned ΔF/F_0_ data were assessed within a 5 s peri-movement window to produce a time series for the decoding accuracy. At each time point, leaving one trial out (test trial), the likelihood of determining push or pull was estimated based on the remaining trials (training set). The leave-one-out procedure was then repeated for all trials to calculate a mean decoding accuracy time series for each neuron. The resulting time series were analyzed within a peri-movement epoch - the peri-movement epoch began at -0.15 s relative to motion index onset and ended based on the peak ΔF/F_0_ response of each neuron; the position of the median peak was calculated for each movement type and the later of these time points used as the cut off. To identify neurons with decoding performance above chance, the bootstrapped distributions of decoding accuracy scores were compared against a threshold value for each session. Only neurons with at least 1 bin significantly higher than threshold were defined as high decoding accuracy. The threshold for each session was calculated based on modeled data composed of random samples from a Gaussian distribution with the same number of trials as the experimental data. For each session, modeled data accuracy was calculated 1000 times, assuming a prior probability of 50:50, and the mean + 2 SD was used as the threshold for significance. For population level classification of movement type, we employed logistic regression. As above, the decoding accuracy of time series for each population was generated via leave-one-out design looped over all the trials in a given session. Population decoding accuracy was defined as the maximum decoding accuracy in any 250 ms bin within the peri-movement epoch. Population decoding was also performed on subsets of the entire population. Neurons were removed from the population one at a time, either in order from highest to lowest decoding accuracy score or randomly, with the network retrained for each iteration. The process was repeated 25 times in the random condition and the median of all responses used as the representative example for comparison with the ordered removal condition. Subpopulations of neurons decoding significantly above chance were determined by comparing decoding scores with a shuffled dataset (sampled randomly from 1000 time points with the trial labels (push or pull) randomized for each sample). If confidence intervals from the population data did not overlap with those of the shuffled data, population scores were deemed to be above chance. In 3/12 FOVs of the layer 5 dataset with low proportions of high decoding neurons and / or low trial numbers, the population decoding accuracy was never significantly above chance. These FOVs were excluded from the comparison between ordered and random removal of neurons.

### Dimensionality reduction

Raw fluorescence traces for all trials with successful movements in a 7.5 s peri-movement window were concatenated, filtered with a three frame (75 ms) wide boxcar kernel, whitened and transformed with principal component analysis. The principal components (PCs) corresponding to the 16 highest eigenvalues, which corresponded to an average 83% (range [77 94]) cumulative explained variance, were analyzed. To compute trajectories in PC space, PC projections for all trials were averaged (separately for push or pull) and the variance and 95% confidence intervals for each time point estimated via 100-fold bootstrapping. The separability of the trajectories for push or pull was computed in each PC separately as *d’(t) =* |*mpush(t) - m*_*pull*_*(t)*| */* √*0*.*5(v*_*push*_*(t) + v*_*pull*_*(t))*, where *m*_*push*_*(t)* and *m*_*pull*_*(t)* are the mean trajectories for push and pull, and v_push_(t) and v_pull_(t) the corresponding variances, estimated from trials. The separability *d’(t)* was bootstrapped from 400 samples, and variance and 95% confidence intervals estimated from this sample. *d’(t)* was computed for all frames from movement onset to completion, where the latter was the longest movement duration recorded in each session. PCs were considered separable if the difference between d’(t) and *d*_*shuffle*_*’(t)* (obtained in the same way from trial-shuffled data) was within the 95% confidence interval, which was estimated from the sum of the relevant bootstrapped variances. For each FOV, the largest significant *d’(t)* was used; in 1/12 FOVs no PCs showed significant separability (this FOV also had a chance level population decoding score), and were excluded in the summary data.

### Spatiotemporal organization

To assess the functional (temporal) organization of simultaneously recorded populations of neurons, pairwise correlation coefficients (Pearson’s r) from the smoothed (1 s LOESS filter) and motion index-aligned ΔF/F_0_ within the peri-movement epoch were compared. Data were split based on their decoding accuracy scores and the bootstrapped median difference between high decoding accuracy neurons and those of the population were subtracted and a median difference calculated per sample. This process was repeated 10,000 times to generate a distribution for high decoding neurons versus the entire population and the same sampling procedure was used to investigate the correlations within low decoding accuracy neurons. To investigate spatial clustering, bootstrapped median differences between high decoding accuracy neurons and the population using pairwise distances (defined as the Euclidean distance between the centroids of manually drawn ROIs from 2-photon imaging processing) were compared. A Generalized Linear Mixed-Effects Model:

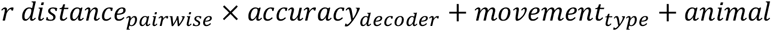

was used to model the pairwise correlation coefficient as a function of pairwise distance (continuous), decoding accuracy and movement type. Pairwise distance and decoding accuracy were modeled as interacting fixed terms, while movement type and animal were modeled as random intercepts to account for the dependency of the measurements on observations from the same animal and across the different movement types. The model was estimated using the restricted maximum likelihood, or REML, method (Bartlett and Fowler, 1937). Model assumptions were verified by comparing residual versus fitted values for each covariate in the model against each covariate removed from the model.

### Statistics

Data analysis was performed using custom-written scripts in MATLAB 2019a or Python 3. Where multiple measurements were made from a single animal, suitable weights were used to evaluate summary population statistics. Statistical significance was considered when p < 0.05 unless otherwise stated. Data were tested for normality and parametric/non-parametric tests were used as appropriate and as detailed in the text. The GLMM was designed in Python using the statsmodels library (Seabold and Perktold, 2010).

