## Supplementary information for "Spatiotemporal organization of movement-invariant and movement-specific signaling in the output layer of motor cortex"

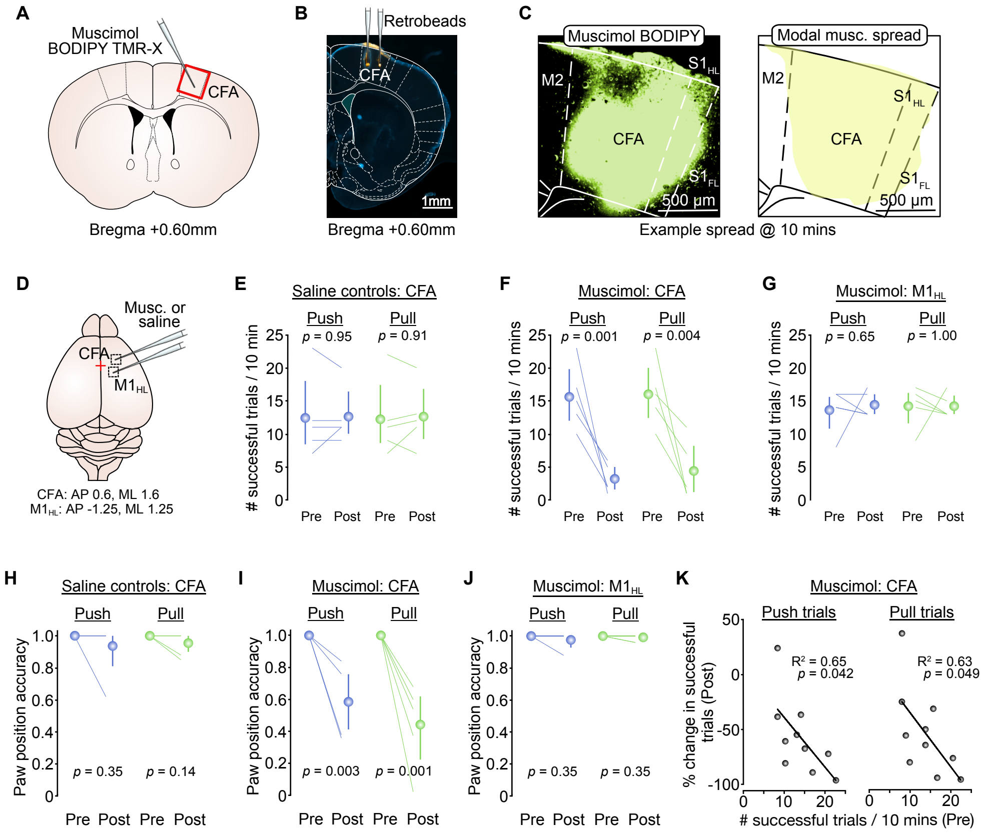


**Figure S1 - Muscimol inactivation of CFA affects forelimb posture and task success.**

(A) Injection of muscimol BODIPY TMR-X into caudal forelimb area (CFA).
(B) Muscimol injection sites visualized using fluorescent green retrobeads.
(C) *Left*, example image of fluorescent muscimol spread in CFA at 10 mins post injection. *Right*, modal spread of fluorescent muscimol in CFA (i.e., area in which fluorescence is present across all mice) (N = 3 mice). M2, secondary motor cortex; $S1_{HL}$, primary hindlimb somatosensory cortex; $S1_{FL}$, primary forelimb somatosensory cortex.
(D) Focal muscimol inactivation of CFA or hindlimb motor cortex ($M1_{HL}$), 0.6 mm anterior, 1.6 mm lateral of bregma and 1.25 mm posterior, 1.25 mm lateral of bregma, respectively. Red cross denotes bregma.
(E-G) Number of successful push (blue) and pull (green) trials in a 10 min period before (Pre) and after (Post) injection of (E) saline into CFA, (F) muscimol into CFA or (G) muscimol into $M1_{HL}$ (N = 5 mice), paired t-test. Colored lines, individual mice. Symbols, population means ± 95% CI.
(H-J) Paw position accuracy at the point of cue presentation before (Pre) and 10 mins after (Post) (H) saline into CFA, (I) muscimol into CFA or (J) muscimol into $M1_{HL}$ (N = 5 mice), paired t-test. Colored lines, individual mice. Symbols, population means ± 95% CI.
(K) Correlation between the number of successful push (left) and pull (right) trials before (Pre) and the % change in successful trials after (Post) muscimol injection (N = 10 mice). Symbols, individual animals, Black line, linear fit to the data (Pearson's *r*).

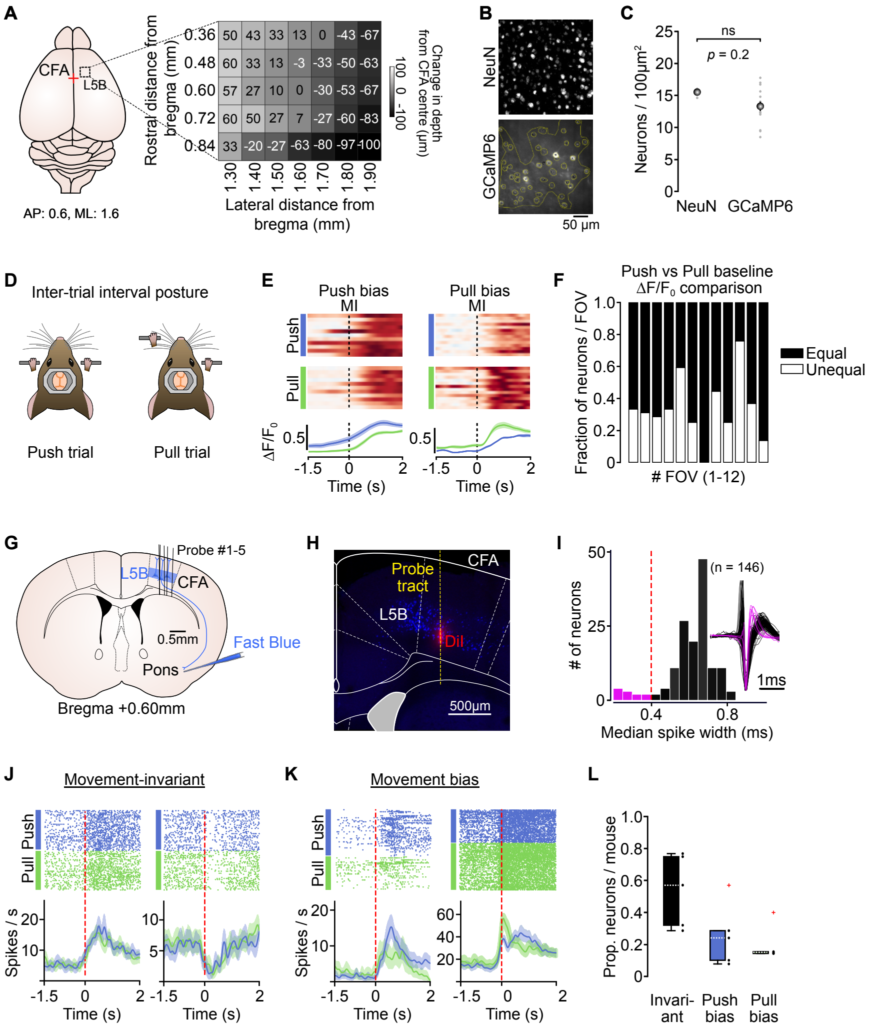


**Figure S2 - Cell density and movement-type classification in layer 5B of CFA.**

(A) *Left*, schematic showing mapped region of caudal forelimb area (CFA) centred on 0.6 mm anterior, 1.6 mm medial of bregma. Red cross denotes bregma; *Right*, heatmap indicating changes in layer 5B depth across a range of cortical coordinates. Values represent the mean depth in µm of the upper boundary of Layer 5B (N = 3 mice).
(B) *Top*, representative image of NeuN stained layer 5B neurons; *Bottom*, representative two-photon image of L5B neurons expressing GCaMP6s. Small circles depict regions-of-interest (yellow) drawn around individual neurons within a larger bounded area excluding blood vessels.
(C) Average number of NeuN versus GCaMP6s expressing neurons / 100µm$^2$ in layer 5B of CFA. Gray dots represent individual slices, bars depict s.e.m. (NeuN, n = 3 slices, N = 1 mouse; GCaMP6s, n = 15 slices, N = 7 mice; two-sample t-test).
(D) Schematic depicting inter-trial posture for push (left) and pull (right) trials.
(E) Activity of two neurons with differences in inter-trial baseline $\Delta F/F_0$. *Left*, example neuron with push bias. *Right*, example neuron with pull bias. *Top*, raster showing normalized $\Delta F/F_0$ across successive push (blue) or pull (green) trials; *Bottom*, mean $\Delta F/F_0$ ± 95% CI for push and pull trials. Dashed lines, movement initiation (MI).
(F) Proportion of neurons per field-of-view (FOV) with equal (black) or unequal (white) inter-trial $\Delta F/F_0$ baselines (n = 486 neurons from 12 FOVs, N = 6 mice).
(G) Silicone probe recordings of putative layer 5B (L5B) neurons in CFA. Approximate position of probes within CFA is depicted. The upper and lower boundaries of layer 5B are defined based on retrograde labeling of pyramidal tract (PT) neurons after FastBlue injection into the pons (N = 5 mice).
(H) Retrograde labeling of pons-projecting PT neurons within CFA (blue) with silicon probe tract visualized using DiI (red). Dashed line, overlay of probe tract.
(I) Histogram of spike durations, highlighting putative interneurons (purple) and pyramidal neurons (black). Inset, trough-aligned mean spike waveforms, normalized based on peak to trough height (n = 148 units, N = 5 mice).
(J) Activity of two example movement-invariant layer 5 CFA neurons. *Top*, Spike rasters during push (blue) and pull (green) trials. Dots represent individual spikes. *Bottom*, peri-stimulus time histograms depicting mean ± s.e.m. firing rate during push (blue) and pull (green) trials. Red dashed lines, movement initiation (MI).
(K) Activity of two example layer 5 CFA neurons with push (left) and pull (right) bias. *Top*, Spike rasters during push (blue) and pull (green) trials. Dots represent individual spikes. *Bottom*, peri-stimulus time histograms depicting mean ± s.e.m. firing rate during push (blue) and pull (green) trials. Red dashed lines, movement initiation (MI).
(L) Proportion of invariant, push- and pull-biased neurons per mouse (n = 71 neurons, N = 5 mice). Black dots represent individual mice. Red crosses, identified outliers.

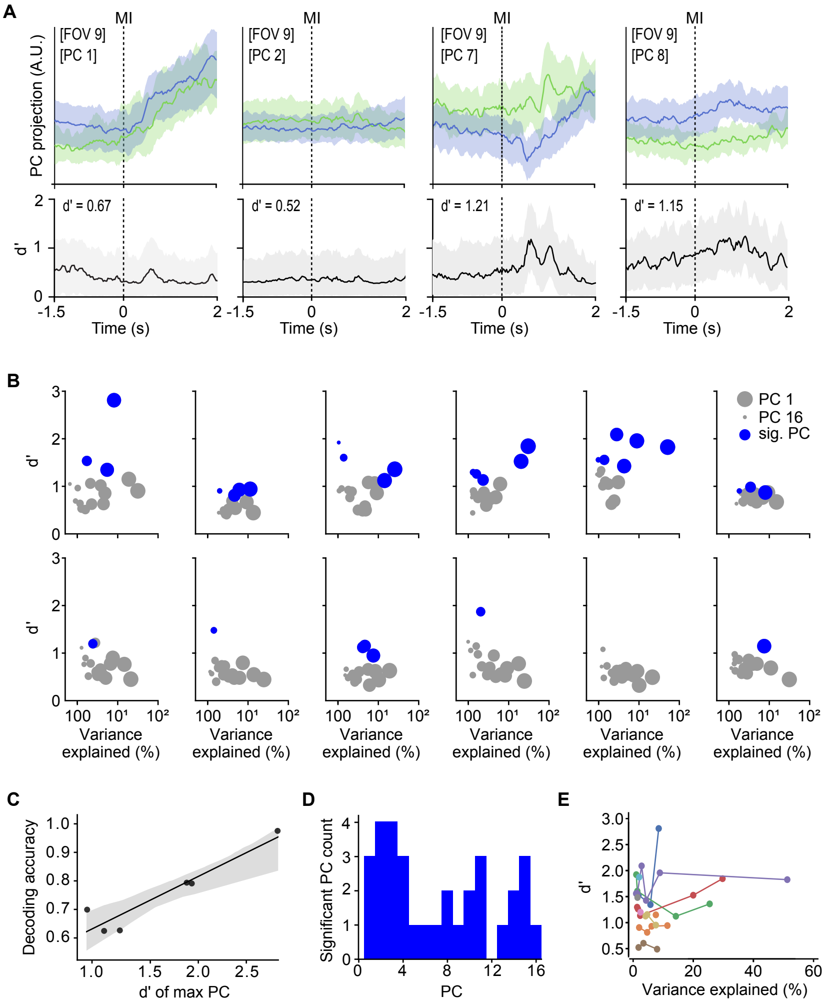

**Figure S3 - Movement type is represented in both leading and higher principal components.**

(A) *Top*, example trajectories of 4 principal components (PC) from a representative field-of-view (FOV) during push (blue) and pull (green trials). *Bottom*, discrimination index (d') calculated from the corresponding PCs. Inset, maximum d' of each PC. Thick lines, mean ± 95% CI. Dashed lines, movement initiation (MI).
(B) Variance explained as a function of discrimination index (d') for all PCs in each FOV. *Top*, FOVs 1-6. *Bottom*, FOVs 7-12. Dot size (large to small) represents PC rank (1-16). Blue dots are significant PCs.
(C) Discrimination index (d') of the highest significant PC vs population decoding accuracy (Figure 3) for each mouse (N = 6 mice). Black line, linear regression fit ± 95% CI.
(D) Histogram of significant PCs across all 12 FOVs.
(E) Significance explained as a function of discrimination index (d') for all significant PCs. Dots represent individual PCs. Lines and colours link PCs from the same FOV.

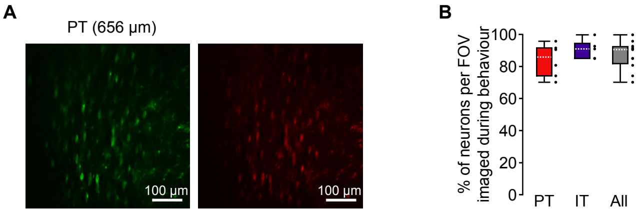


**Figure S4 - Quantifying proportions of imaged neurons per field-of-view.**

(A) Example field-of-view (FOV) showing PT neurons expressing GCaMP6s (left; green) and mRuby (right; red).
(B) Percentage of mRuby positive PT and IT neurons per FOV imaged ($\Delta F/F_0$) during behaviour. Box-and-whisker plots showing median, interquartile range and range. PT (red), IT (purple) and all neurons (gray). Black dots represent individual FOVs. N = 6, 5 and 11, respectively.

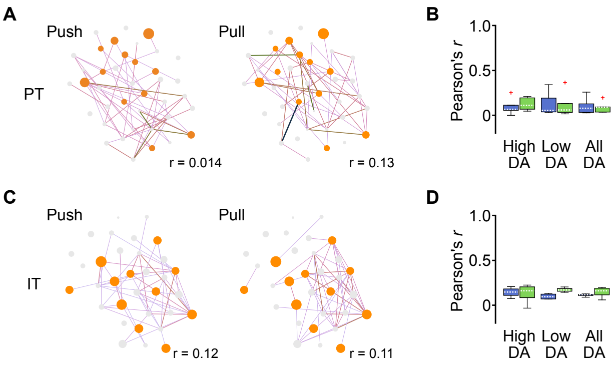


**Figure S5 - Temporal organization of high decoding accuracy neurons in layer 5B during push and pull movements.**

(A) *Left & right*, functional networks constructed from the pairwise activity correlations from a representative PT field-of-view (FOV) for push (left) or pull (right) trials. Line color (light to dark) and width correspond to increasing values of Pearson's *r*. Neurons are plotted as nodes in Euclidean space with color and size relating to increasing decoding accuracy.
(B) Box-and-whisker plots showing the median, interquartile range and range of correlation coefficients across mice for high decoding accuracy (HDA), low decoding accuracy (LDA) and all neurons. Data are shown for both push (blue) and pull (green) trials. Red crosses, identified outliers.
(C) *Left & right*, functional networks constructed from the pairwise activity correlations from a representative IT FOV for push (left) or pull (right) trials. Line color (light to dark) and width correspond to increasing values of Pearson's *r*. Neurons are plotted as nodes in Euclidean space with color and size relating to increasing decoding accuracy.
(D) Box-and-whisker plots showing the median, interquartile range and range of correlation coefficients across mice for high decoding accuracy (HDA), low decoding accuracy (LDA) and all neurons. Data are shown for both push (blue) and pull (green) trials.

**Table of results**

$\bar{X}$ = mean $\tilde{X}$ = median SD = standard deviation IQR = interquartile range
fov = fields-of-view H / LDA = high / low decoding accuracy GLMM = generalised linear mixed-effects model

| Figure | Description | Sample size | Result | Variance | Confidence intervals (95%) | Statistical test result |
|---|---|---|---|---|---|---|
| 1C-E | Training time (days) | N = 24 | $\tilde{X}$ = 10.5 | IQR = 4 | | |
| 1C | Reaction time (s; push / pull) | N = 24 | $\tilde{X}$ = 0.132 / 0.135 | IQR = 0.058 / 0.043 | | |
| | Movement duration (s; push / pull) | N = 24 | $\tilde{X}$ = 706 / 422 | IQR = 352 / 438 | | |
| 1D | Successful trials / session (push / pull) | N = 24 | $\tilde{X}$ = 44.5 / 45 | IQR = 9.5 / 8.5 | | |
| 1E | Task success (%; push / pull) | N = 24 | $\tilde{X}$ = 68.0 / 74.5 | IQR = 35.9 / 43.4 | | |
| 1G | Successful trials / 10min (push muscimol CFA pre / post) | N = 10 | $\bar{X}$ = 13.9 / 5 | SD = 5.22 / 2.94 | [11.1 17.1] / [3.3 6.8] | t(18) = 4.70, *P* = $1.79 \times 10^{-4}$, paired t-test |
| | Successful trials / 10min (pull muscimol CFA pre / post) | N = 10 | $\bar{X}$ = 14.0 / 5.3 | SD = 5.33 / 3.62 | [10.9 17.2] / [3.2 7.4] | t(18) = 4.27, *P* = $4.64 \times 10^{-4}$, paired t-test |
| 1H | Paw position accuracy (push muscimol CFA pre / post) | N = 10 | $\bar{X}$ = 1 / 0.66 | SD = 0 / 0.2 | [1 1] / [0.54 0.77] | t(18) = 5.56, *P* = $2.79 \times 10^{-5}$, paired t-test |
| | Paw position accuracy (pull muscimol CFA pre / post) | N = 10 | $\bar{X}$ = 0.98 / 0.50 | SD = 0.06 / 0.21 | [0.95 1] / [0.36 0.62] | t(18) = 6.85, *P* = $2.07 \times 10^{-6}$, paired t-test |
| S1. E | Successful trials / 10min (push saline CFA pre / post) | N = 5 | $\bar{X}$ = 12.4 / 12.6 | SD = 6.23 / 4.28 | [8.6 18] / [10 16.4] | t(8) = -0.06, *P* = 0.95, paired t-test |

| | | | | | | |
|---|---|---|---|---|---|---|
| | Successful trials / 10min (pull saline CFA pre / post) | N = 5 | $\bar{X}$ = 12.2 / 12.6 | SD = 5.81 / 4.72 | [8.6 17.4] / [9 16.6] | t(8) = -0.12, *P* = 0.91, paired t-test |
| S1. F | Successful trials / 10min (push muscimol CFA pre / post) | N = 5 | $\bar{X}$ = 15.6 / 3.2 | SD = 4.88 / 2.17 | [12.0 19.8] / [1.6 4.8] | t(8) = 5.19, *P* = $8.29 \times 10^{-4}$, paired t-test |
| | Successful trials / 10min (pull muscimol CFA pre / post) | N = 5 | $\bar{X}$ = 16.0 / 4.4 | SD = 4.74 / 4.45 | [12.4 20.0] / [1.2 8.2] | t(8) = 3.99, *P* = $4.0 \times 10^{-3}$, paired t-test |
| S1. G | Successful trials / 10min (push muscimol $M1_{HL}$ pre / post) | N = 5 | $\bar{X}$ = 13.6 / 14.4 | SD = 3.29 / 1.95 | [10.8 15.6] / [13.0 16.0] | t(8) = -0.47, *P* = 0.65, paired t-test |
| | Successful trials / 10min (pull muscimol $M1_{HL}$ pre / post) | N = 5 | $\bar{X}$ = 14.2 / 14.2 | SD = 3.11 / 1.79 | [11.4 16.2] / [13.0 15.8] | t(8) = 0, *P* = 1.0, paired t-test |
| S1. H | Paw position accuracy (push saline CFA pre / post) | N = 5 | $\bar{X}$ = 1 / 0.92 | SD = 0 / 0.17 | [1 1] / [0.77 1] | t(8) = 1.0, *P* = 0.35, paired t-test |
| | Paw position accuracy (pull saline CFA pre / post) | N = 5 | $\bar{X}$ = 1 / 0.95 | SD = 0 / 0.07 | [1 1] / [0.89 1] | t(8) = 1.62, *P* = 0.15, paired t-test |
| S1. I | Paw position accuracy (push muscimol CFA pre / post) | N = 5 | $\bar{X}$ = 1 / 0.59 | SD = 0 / 0.22 | [1 1] / [0.41 0.76] | t(8) = 4.27, *P* = $2.7 \times 10^{-3}$, paired t-test |
| | Paw position accuracy (pull muscimol CFA pre / post) | N = 5 | $\bar{X}$ = 1 / 0.44 | SD = 0 / 0.26 | [1 1] / [0.23 0.62] | t(8) = 4.79, *P* = $1.4 \times 10^{-3}$, paired t-test |
| S1. J | Paw position accuracy (push muscimol $M1_{HL}$ pre / post) | N = 5 | $\bar{X}$ = 1 / 0.98 | SD = 0 / 0.05 | [1 1] / [0.93 1] | 1(8) = 1.0, *P* = 0.35, paired t-test |
| | Paw position accuracy (pull muscimol $M1_{HL}$ pre / post) | N = 5 | $\bar{X}$ = 1 / 0.99 | SD = 0 / 0.02 | [1 1] / [0.98 1] | 1(8) = 1.0, *P* = 0.35, paired t-test |
| S1. K | Successful trial vs % change in successful trials (push / pull) | N = 10 | | | | $r^2$ = -0.65 *P* = 0.042 / $r^2$ = -0.63, *P* = 0.049 |

| | | | | | | |
|---|---|---|---|---|---|---|
| 2D | Prop. of responsive neurons (non / movement / reward) | N = 6<br>fov = 12<br>cell = 653 | $\bar{X}$ = 20.9 / 73.5 / 5.7 | SD = 15.1 / 16.0 / 3.7 | [12.8 29.1] / [64.7 81.8] / [3.7 7.7] | |
| 2E | Trial-trial correlation - ΔF/F0 vs motion index for push trials (motion index correlation 0.1 / 0.3 / 0.5 0.7) | N = 6<br>fov = 12<br>trials = 2931 | $\tilde{X}$ = 0.36 / 0.62 / 0.75 / 0.80 | | [0.24 0.40] / [0.46 0.68] / [0.60 0.86] / [0.65 0.89] | |
| | Trial-trial correlation - ΔF/F0 vs motion index for pull trials (motion index correlation 0.1 / 0.3 / 0.5 0.7) | N = 6<br>fov = 12<br>trials = 2931 | $\tilde{X}$ = 0.25 / 0.47 / 0.59 / 0.79 | | [0.23 0.27] / [0.37 0.87] / [0.52 0.68] / [0.65 0.89] | |
| 2G | # of movement-responsive neurons (%; bias / invariant) | N = 6<br>fov = 12<br>Cell = 468 | 181 (38.7) / 287 (61.3) | | | |
| 2J | # of movement bias neurons (%; type 1 / 2 / 3 / 4) | N = 6<br>fov = 12<br>Cell = 181 | 136 (75.1) / 25 (13.8) / 15 (8.3) / 5 (2.8) | | | |
| 2L | Average ΔF/F0 trial-trial correlation for push trials (non / push / push) | N = 6<br>fov = 12<br>cell = 468 | $\tilde{X}$ = 0.37 / 0.49 / 0.45 | IQR = 0.37 / 0.47 / 0.39 | [0.12 0.56] / [0.21 0.72] / [0.04 0.70] | |
| | Average ΔF/F0 trial-trial correlation for pull trials (non / push / push) | N = 6<br>fov = 12<br>cell = 468 | $\tilde{X}$ = 0.31 / 0.30 / 0.53 | IQR = 0.25 / 0.32 / 0.33 | [0.10 0.65] / [0.08 0.75] / [0.06 0.61] | |
| 2M | Prop. of movement-responsive neurons (non / push / pull) | N = 6<br>fov = 12<br>cell = 468 | $\tilde{X}$ = 59.8 / 14.3 / 11.8 | IQR = 31.4 / 15.9 / 19.5 | | |
| S2. C | Neurons / 100 μm$^2$ (NeuN / GCaMP6s) | N = 1 / 7<br>fov = 3 / 15 | $\bar{X}$ = 15.6 / 13.3 | SD = 0.9 / 2.7 | | t(16) = 1.75, *P* = 0.2, Student's t-test |
| S2. E-F | Prop. of biased neurons (equal / unequal baseline) | N = 6<br>fov = 12 | $\tilde{X}$ = 69.5 / 30.5 | IQR = 8.2 / 8.2 | | |

| | | | | | | |
|---|---|---|---|---|---|---|
| | | cell = 181 | | | | |
| S2. G-L | # units (%; pyramidal / interneurons) | N = 5<br>n = 146 | 137 (93.8) / 9 (6.2) | | | |
| | # units (%; non / move / reward) | N = 5<br>n = 137 | 61 (45.2) / 72 (52.6) / 4 (2.9) | | | |
| S2. L | Prop. of movement-responsive neurons (non / push / pull) | N = 5<br>n = 72 | $\tilde{X}$ = 0.57 / 0.15 / 0.24 | IQR = 0.43 / 0.01 / 0.19 | | |
| 3A | Prop. of movement-responsive neurons (LDA / HDA) | N = 6<br>fov = 12<br>cell = 468 | 0.63 / 0.37 | | | |
| 3B | Decoding accuracy (single cell / population) | N = 6<br>fov = 12 | $\tilde{X}$ = 0.61 / 0.75 | IQR = 0.07 / 0.16 | | W = 1, Z = -2.20, $P = 2.8 \times 10^{-2}$, Wilcoxon signed rank test |
| 3F | Prop. neurons removed (high-low / random) | N = 6<br>fov = 9 | $\tilde{X}$ = 0.21 / 0.64 | IQR = 0.5 / 0.57 | | W = 1, Z = -2.20, $P = 2.8 \times 10^{-2}$, Wilcoxon signed rank test |
| S3. C | Population decoding vs max d’ | N = 6<br>fov = 10 | | | | $r^2$ =0.88, F(1,5) = 29.95, $P = 5.4 \times 10^{-3}$ |
| 4E | # of movement-responsive neurons PT (%; bias / invariant) | N = 5<br>fov = 6<br>cell = 171 | 46 (26.9) / 125 (73.1) | | | |
| | # of movement bias neurons PT (%; type 1 / 2 / 3 / 4) | N = 5<br>fov = 6<br>cell = 46 | 36 (78.2) / 9 (19.6) / 1 (2.2) / 0 (0) | | | |
| | Prop. of movement-responsive neurons PT (non / push / pull) | N = 5<br>fov = 6<br>cell = 171 | $\tilde{X}$ = 0.75 / 0.10 / 0.14 | IQR = 0.21 / 0.19 / 0.11 | | |

| | | | | | | |
|---|---|---|---|---|---|---|
| 4G | # of movement-responsive neurons IT (%; bias / invariant) | N = 4<br>fov = 5<br>cell = 110 | 54 (49.1) / 56 (50.9) | | | |
| | # of movement bias neurons IT (%; type 1 / 2 / 3 / 4) | N = 4<br>fov = 5<br>cell = 54 | 34 (62.9) / 15 (27.8) / 2 (3.7) / 3 (5.6) | | | |
| | Prop. of movement-responsive neurons IT (non / push / pull) | N = 4<br>fov = 5<br>cell = 110 | $\tilde{X}$ = 0.51 / 0.21 / 0.32 | IQR = 0.11 / 0.26 / 0.31 | | |
| 4I | Decoding accuracy PT (single / population) (HDA only) | N = 5<br>fov = 6<br>cell = 58 | $\tilde{X}$ = 0.55 / 0.68 | IQR = 0.02 / 0.14 | | t(4) = -3.04, *P* = 0.04 |
| | Decoding accuracy IT (single / population) (HDA only) | N = 4<br>fov = 5<br>cell = 43 | $\tilde{X}$ = 0.56 / 0.73 | IQR = 0.04 / 0.21 | | t(3) = -2.95, *P* = 0.06 |
| | Decoding accuracy population (PT / IT) (HDA only) | N = 5 / 4<br>fov = 6 / 5<br>cell = 58 / 43 | $\tilde{X}$ = 0.68 / 0.73 | IQR = 0.14 / 0.21 | | t(7) = 0.40, *P* = 0.70 |
| S4. B | % of all neurons per FOV imaged during behaviour (PT / IT / All) | N = 5 / 4 / 9<br>fov = 6 / 5 / 11 | $\tilde{X}$ = 85.8 / 91.2 / 90.7 | IQR = 17.4 / 9.3 / 10.7 | | |
| 5B | Norm. prop. $\Delta F/F_0$ onsets push PT (ms; HDA / LDA). ANOVA [animal:cell type: onset time] | N = 5<br>fov = 6<br>cell = 238 | $\tilde{X}$ = 210 / 113 | IQR = 709 / 662 | | F(4) = 0, *P* = 1 / F(1) = 0, *P* = 1 / F(17) = 5.12, *P* = $8.32 \times 10^{-9}$ |
| | Norm. prop. $\Delta F/F_0$ onsets pull PT (ms; HDA / LDA). ANOVA [animal:cell type: onset time] | N = 5<br>fov = 6<br>cell = 238 | $\tilde{X}$ = 145 / 177 | IQR = 226 / 500 | | F(3) = 0, *P* = 1 / F(1) = 0, *P* = 1 / F(17) = 8.57, *P* = $3.35 \times 10^{-15}$ |

| | | | | | | |
|---|---|---|---|---|---|---|
| | Norm. prop. $\Delta F/F_0$ onsets push IT (ms; HDA / LDA). ANOVA [animal:cell type: onset time] | N = 4<br>fov = 5<br>cell = 137 | $\tilde{X}$ = 500 / 339 | IQR = 524 / 677 | | F(3) = $1.1\times10^{-14}$, *P* = 1 / F(1) = 0, *P* = 1 / F(17) = 7.31, *P* = $4.85\times10^{-12}$ |
| | Norm. prop. $\Delta F/F_0$ onsets pull IT (ms; HDA / LDA). ANOVA [animal:cell type: onset time] | N = 4<br>fov = 5<br>cell = 137 | $\tilde{X}$ = 403 / 468 | IQR = 444 / 710 | | F(3) = 0, *P* = 1 / F(1) = 0, *P* = 1 / F(17) = 3.81, *P* = $6.85\times10^{-6}$ |
| 5D | Pairwise correlation coefficient PT (r; push; HDA vs ALL / LDA vs ALL) | N = 5<br>fov = 6<br>cell = 171 | HDA $\tilde{X}$ = 0.07<br>LDA $\tilde{X}$ = 0.06<br>ALL $\tilde{X}$ = 0.08 | HDA IQR = 0.05<br>LDA IQR = 0.16<br>ALL IQR = 0.10 | [-0.103 0.087] / [-0.085 0.094] | *P* = 0.84 / 1.0 |
| | Pairwise correlation coefficient IT (r; push; HDA vs ALL / LDA vs ALL) | N = 4<br>fov = 5<br>cell = 110 | HDA $\tilde{X}$ = 0.15<br>LDA $\tilde{X}$ = 0.09<br>ALL $\tilde{X}$ = 0.12 | HDA IQR = 0.07<br>LDA IQR = 0.05<br>ALL IQR = 0.01 | [-0.093 0.081] / [-0.073 0.076] | *P* = 1.0 / 0.96 |
| 5E | Pairwise corr. Vs distance PT (HDA / LDA) | N = 5<br>fov = 6<br>cell = 171 | Spearman r = -0.1 / 0.0 | | | *P* = 0.87 / 1.0 |
| | GLMM PT<br>r distance pairwise accuracydecoder + movementtype+animal | N = 5, n = 3024 observations | | | | *P* = 0.33 |
| | Pairwise corr. Vs distance IT (HDA / LDA) | N = 4<br>fov = 5<br>cell = 110 | Spearman r = -0.40 / 0.0 | | | *P* = 0.6 / 1.0 |
| | GLMM IT<br>r distance pairwise accuracydecoder + movementtype+animal | N = 5, n = 1562 observations | | | | *P* = 0.49 |
| 5G | Pairwise distance PT (µm; HDA vs ALL / LDA vs ALL) | N = 5<br>fov = 6<br>cell = 171 | HDA $\tilde{X}$ = 226<br>LDA $\tilde{X}$ = 273 | HDA IQR = 26<br>LDA IQR = 60<br>ALL IQR = 53 | [-69 102] / [-62 69] | *P* = 1.0 / 1.0 |

| | | | | | | |
|---|---|---|---|---|---|---|
| | | | ALL $\tilde{X}$ = 264 | | | |
| | Pairwise distance IT (μm; HDA vs ALL / LDA vs ALL) | N = 4<br>fov = 5<br>cell = 110 | HDA $\tilde{X}$ = 228<br>LDA $\tilde{X}$ = 220<br>ALL $\tilde{X}$ = 229 | HDA IQR = 14<br>LDA IQR = 56<br>ALL IQR = 58 | [-82 67] / [-63 68] | *P* = 0.88 / 1.0 |
| S5. B | Pairwise correlation coefficient PT (r; pull; HDA vs ALL / LDA vs ALL) | N = 5<br>fov = 6<br>cell = 171 | HDA $\tilde{X}$ = 0.11<br>LDA $\tilde{X}$ = 0.06<br>ALL $\tilde{X}$ = 0.09 | HDA IQR = 0.12<br>LDA IQR = 0.10<br>ALL IQR = 0.06 | [-0.1 0.15] / [-0.1 0.09] | *P* = 1.17 / 0.79 |
| S5. D | Pairwise correlation coefficient IT (r; pull; HDA vs ALL / LDA vs ALL) | N = 4<br>fov = 5<br>cell = 110 | HDA $\tilde{X}$ = 0.16<br>LDA $\tilde{X}$ = 0.17<br>ALL $\tilde{X}$ = 0.16 | HDA IQR = 0.12<br>LDA IQR = 0.03<br>ALL IQR = 0.08 | [-0.08 0.07] / [-0.07 0.1] | *P* = 0.89 / 1.28 |
